# Automatic Spatial Estimation of White Matter Hyperintensities Evolution in Brain MRI using Disease Evolution Predictor Deep Neural Networks

**DOI:** 10.1101/738641

**Authors:** Muhammad Febrian Rachmadi, Maria del C. Valdés-Hernández, Stephen Makin, Joanna Wardlaw, Taku Komura

## Abstract

Previous studies have indicated that white matter hyperintensities (WMH), the main radiological feature of small vessel disease, may evolve (i.e., shrink, grow) or stay stable over a period of time. Predicting these changes are challenging because it involves some unknown clinical risk factors that leads to a non-deterministic prediction task. In this study, we propose a deep learning model to predict the evolution of WMH from baseline to follow-up (i.e., 1-year later), namely “Disease Evolution Predictor” (DEP) model, which can be adjusted to become a non-deterministic model. The DEP model receives a baseline image as input and produces a map called “Disease Evolution Map” (DEM), which represents the evolution of WMH from baseline to follow-up. Two DEP models are proposed, namely DEP-UResNet and DEP-GAN, which are representatives of the supervised (i.e., need expert-generated manual labels to generate the output) and unsupervised (i.e., do not require manual labels produced by experts) deep learning algorithms respectively. To simulate the non-deterministic and unknown parameters involved in WMH evolution, we modulate a Gaussian noise array to the DEP model as auxiliary input. This forces the DEP model to imitate a wider spectrum of alternatives in the prediction results. The alternatives of using other types of auxiliary input instead, such as baseline WMH and stroke lesion loads are also proposed and tested. Based on our experiments, the fully supervised machine learning scheme DEP-UResNet regularly performed better than the DEP-GAN which works in principle without using any expert-generated label (i.e., unsupervised). However, a semi-supervised DEP-GAN model, which uses probability maps produced by a supervised segmentation method in the learning process, yielded similar performances to the DEP-UResNet and performed best in the clinical evaluation. Furthermore, an ablation study showed that an auxiliary input, especially the Gaussian noise, improved the performance of DEP models compared to DEP models that lacked the auxiliary input regardless of the model’s architecture. To the best of our knowledge, this is the first extensive study on modelling WMH evolution using deep learning algorithms, which deals with the non-deterministic nature of WMH evolution.

## 1. Introduction

White matter hyperintensities (WMH), together with lacunar ischaemic strokes, lacunes, cerebral microbleeds, and perivascular spaces, are the main neuroradiological features of cerebral small vessel disease (SVD) (Wardlaw et al., 2013). WMH can be observed in T2-weighted and T2-fluid attenuated inversion recovery (T2-FLAIR) brain magnetic resonance images (MRI), sharing similar neuroradiological characteristics as the lacunar ischaemic infarcts and enlarged perivascular spaces (del C. Valdés Hernández et al., 2013). Clinically, WMH have been associated with stroke, ageing, and dementia progression (Prins and Scheltens, 2015; Wardlaw et al., 2017a). Recent studies have shown that WMH may decrease (i.e., shrink/regress), stay unchanged (i.e., stable), or increase (i.e., grow/progress) over a period of time (Ramirez et al., 2016). Variations in the WMH burden over time have been associated with patients’ comorbidities and clinical outcome (Chappell et al., 2017; Wardlaw et al., 2017b). In this study, we refer to theses changes as “evolution of WMH”.

Predicting the evolution of WMH is challenging because the rate and direction of WMH evolution varies considerably across studies (Schmidt et al., 2016; van Leijsen et al., 2017a,b) and several risk factors, either commonly or not fully known, could be involved in their progression (Wardlaw et al., 2017b). For example, some risk factors and predictors that have been commonly associated with WMH progression are baseline WMH volume (Schmidt et al., 2003; Sachdev et al., 2007; van Dijk et al., 2008; Wardlaw et al., 2017b; Chappell et al., 2017), blood pressure or hypertension (Veldink et al., 1998; Schmidt et al., 2002b; van Dijk et al., 2008; Godin et al., 2011; Verhaaren et al., 2013), age (van Dijk et al., 2008), current smoking status (Power C et al., 2015), previous stroke and diabetes (Gouw et al., 2008; Wardlaw et al., 2017b), and genetic properties (Schmidt et al., 2002a, 2011; Godin et al., 2009; Luo et al., 2017). Surrounding regions of WMH that may appear like normal appearing white matter (NAWM) with less structural integrity, usually called the “penumbra of WMH” (Maillard et al., 2011), have also been reported as having a high risk of becoming WMH over time (Maillard et al., 2014; Pasi et al., 2016). On the other hand, regression of WMH volume has been reported in several radiological observations on MRI, such as after cerebral infraction (Moriya et al., 2009), strokes (Durand-Birchenall et al., 2012; Cho et al., 2015; Wardlaw et al., 2017b), improved hepatic encephalopathy (Mínguez et al., 2007), lower blood pressure (Wardlaw et al., 2017b), liver transplantation (Rovira Cañellas et al., 2007), and carotid artery stenting (Yamada et al., 2010). While a recent study suggested that areas of shrinking WMH were actually still damaged (Jiaerken et al., 2018), a more recent study showed that WMH regression did not accompany brain atrophy and suggested that WMH regression follows a relatively benign clinical course (van Leijsen et al., 2019).

In this study, we propose an end-to-end training model for automatically predicting and spatially estimating the dynamic evolution of WMH from *baseline* to the *following time point* using deep neural networks called “Disease Evolution Predictor” (DEP) model (discussed in Section 2.2). The DEP model produces a map named “Disease Evolution Map” (DEM) which characterises each WMH or brain tissue voxel as progressing, regressing, or stable WMH (discussed in Section 2.1). For this study we have chosen deep neural networks due to their exceptional performance on WMH segmentation (Rachmadi et al., 2017; Li et al., 2018; Kuijf et al., 2019). We use a Generative Adversarial Network (GAN) (Goodfellow et al., 2014) and the U-Residual Network (UResNet) (Guerrero et al., 2018) as base architectures for the DEP model. These architectures represent the *state-of-the-art* deep neural network models. In other words, GANs do not need expert-annotated manual labels in the learning process as they learn the regularities within the input data to estimate the unknown patterns without the need of pre-existing labels (i.e., unsupervised), whilst UResNet adjusts each layer’s weights by recurrently optimising its response in a regularisation process that needs a model expert-annotated label to compare against (i.e., supervised).

This study differs from previous studies on predictive modelling in the fact that we are interested in predicting the evolution of specific neuroradiological MRI features (i.e., WMH in T2-FLAIR), not the progression of a disease as a whole and/or its effect. For example, previous studies have proposed methods for predicting the progression from mild cognitive impairment to Alzheimer’s disease (Spasov et al., 2019; Korolev et al., 2016; Hinrichs et al., 2011) and progression of cognitive decline in Alzheimer’s disease patients (Choi et al., 2018). These studies have used multiple kernel learning classification approaches, which can incorporate non-imaging inputs to increase specificity in the classification, leading to relative accurate prediction of transitional stages or classes. However, it is unclear whether such approaches can cope with voxel-based prediction amidst the ill-posed boundary conditions that distinguish normal from abnormal tissue (i.e., WMH) in diseases that have a wide range of structural abnormalities of different degrees coexisting simultaneously, like the case of small vessel disease as previously mentioned.

Our proposed DEP model generates three outcomes: 1) prediction of WMH volumetric changes (i.e., either progressing or regressing), 2) estimation of WMH spatial changes, and 3) spatial distribution of white matter evolution at the voxel-level precision. Thus, using the DEP model, clinicians can estimate the size, extent, and location of WMH in time to study their progression/regression in relation to clinical health and disease indicators, for ultimately design more effective therapeutic interventions (Rachmadi et al., 2019). Results and evaluations can be seen in Section 4.

To the best of our knowledge, this is the first extensive study on modelling the dynamic change and evolution of WMH, especially using deep learning algorithms. Some relevant studies which use non-deep learning algorithms for predicting other types of brain lesions have been previously proposed. For example, a study that modelled ischemic stroke lesion dynamics using longitudinal metamorphosis (Rekik et al., 2014) and a study that modelled low-grade gliomas using tumor growth parameters estimation (Rekik et al., 2013). Most of these studies use mathematical models that consider a limited number of clusters with well-defined boundaries, which is not the case of the WMH (Rekik et al., 2012; Elazab et al., 2018). However, some recent relevant studies have proposed the use of deep neural networks for estimating the brain tumor growth’s parameters (Ezhov et al., 2019) with emphasis in the dynamic of glioma growth (Petersen et al., 2019). The difference between our study and these recent studies is that our proposed DEP model allows modulating non-image information using auxiliary input (discussed in Section 2.3).

This is an extensive study which expands our previous work in MICCAI 2019 (Rachmadi et al., 2019) where the first study on probabilistic prediction method for WMH evolution was proposed. The main contributions of this study, not addressed in our previous work are as follows.

1. We propose and evaluate the use of three different input modalities for the DEM: 1) irregularity map (IM) (Rachmadi et al., 2019), 2) probability map (PM) generated from a supervised deep learning WMH segmentation method, and 3) binary WMH label (LBL) generated by an expert or highly trained analyst.
2. We performed an ablation study of using different GAN architectures for DEP-GAN model, namely 1) Wasserstein GAN with gradient penalty (WGAN-GP), 2) visual attribution GAN (VA-GAN), 3) DEP-GAN with 1 critic (DEP-GAN-1C), and 4) DEP-GAN with 2 critics (DEP-GAN-2C).
3. We investigated three different levels of human supervision in predicting WMH evolution: 1) supervised DEP-UResNet using expert-generated manual labels, 2) unsupervised DEP-GAN using IM produced by an unsupervised segmentation method of LOTSIM (Rachmadi et al., 2019), and 3) semi-supervised DEP-GAN using PM produced by a supervised segmentation method of UResNet.
4. We performed an ablation study of four different types of auxiliary input for DEP model: 1) no auxiliary input, 2) baseline WMH load, 3) baseline WMH and stroke lesions (SL) loads, and 4) Gaussian noise.
5. We performed clinical plausibility analysis of the application of each DEP model in predicting WMH volumetric changes accounting for risk factors of WMH evolution using analysis of covariance (ANCOVA).

## 2. Proposed Methods

### 2.1. Disease Evolution Map (DEM)

To produce a standard representation of WMH evolution, a simple subtraction operation between two irregularity maps from two time points (i.e., baseline assessment from follow-up assessment) named “Disease Evolution Map” (DEM) was proposed in our previous work (Rachmadi et al., 2019). In the present study, we evaluate the use of three different modalities in the subtraction operation: irregularity map (i.e. as per (Rachmadi et al., 2019)), probability map, and binary WMH label.

Irregularity map (IM) is a map/image that describes the “irregularity” level of each voxel with respect to the normal brain tissue using real values between 0 and 1 (Rachmadi et al., 2018b). The IM is unique as it retains some of the original MRI textures (e.g., from the T2-FLAIR image intensities), including gradients of WMH. IM is also independent from a human rater or training data, as it is produced using an unsupervised method (i.e., LOTS-IM) (Rachmadi et al., 2020). Furthermore, previous studies have shown that IM can also be used for WMH segmentation (Rachmadi et al., 2018b), data augmentation of supervised WMH segmentation (Jeong et al., 2019), and simulation of WMH progression and regression (Rachmadi et al., 2018c). DEM resulted from the subtraction of two IMs has values ranging from −1 to 1 (first row of Figure 1). Note how both regression and progression (i.e. blue for negative values and red for positive values) are well represented at the voxel level precision on the DEM obtained from IMs.

**Figure 1:**
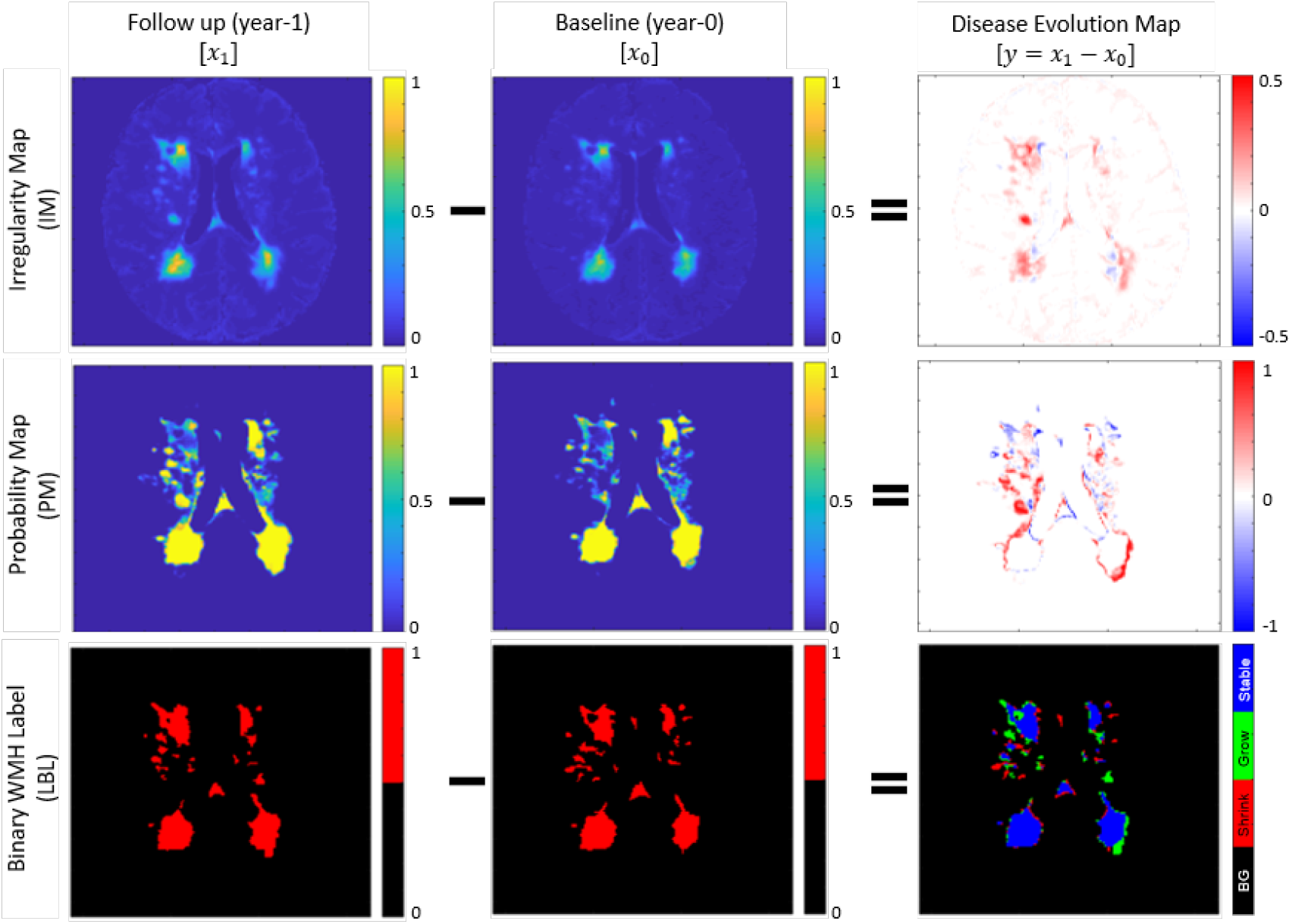
“Disease evolution map” (DEM) (**right**) is produced by subtracting baseline images (**middle**) from follow-up image (**left**). In DEM produced by irregularity map (IM) (**first row**) and probability map (PM) (**second row**), bright yellow pixels represent positive values (i.e., progression) while dark blue pixels represent negative values (i.e., regression). On the other hand, DEM produced by binary WMH label (LBL) (**third row**) has three foreground labels which represent progression or “Grow” (green), regression or “Shrink” (red), and “Stable” (blue). We named this special DEM as three-class DEM label (LBL-DEM).

Probability map (PM) in the present study refers to the WMH segmentation output from a supervised machine learning method. Similar to IM, PM has real values between 0 and 1 which describe the probability for each voxel of being WMH. However, PM differs from IM in the fact that PM only has WMH gradients on the borders of WMH (note that the centres of (big) WMH clusters mostly have probability of 1). Thus, the DEM produced from the subtraction of two PMs also has values ranging from −1 to 1 representing regression and progression respectively, but these are usually located on the WMH clusters’ borders and/or representing small WMH. On the other hand, the rest of DEM’s regions (i.e., the centers of big WMH and non-WMH regions) have value of 0 (see the second row of Figure 1).

Lastly, binary WMH label (LBL) refers to the WMH label produced by an expert’s manual segmentation, which is often considered as gold standard (Valdés Hernández et al., 2015). DEM from LBL can be produced by subtracting the baseline LBL from the follow-up LBL, and each voxel of the resulted image is then labelled as either “Shrink” if it has value below zero, “Grow” if it has value above zero, or “Stable” if it has value of zero. We refer this DEM as three-class DEM label (LBL-DEM), and its depiction can be seen in the bottom-right of Figure 1.

### 2.2. Disease Evolution Predictor (DEP) Model using Deep Neural Networks

In this study, two Disease Evolution Predictor (DEP) models are proposed and evaluated: 1) DEP model based on generative adversarial networks (DEP-GAN) (Rachmadi et al., 2019) and 2) DEP model based on UResNet (DEP-UResNet). The differences between DEP-GAN and DEP-UResNet are input/output modalities and level of human supervision involved in the training. For input/output modalities, DEP-GAN uses either IM or PM for both input and output modalities to represent WMH whereas DEP-UResNet uses T2-FLAIR and expert generated three-class DEM label (LBL-DEM) for input and output respectively. In terms of the level of human supervision, DEP-GAN using IM is categorised as unsupervised because the input modality (i.e., IM) is produced by an unsupervised method (i.e., LOTS-IM), DEP-GAN using PM is categorised as semi-supervised because the PM is produced by a supervised deep learning algorithm (i.e., UResNet, see Section 3.2), and DEP-UResNet is categorised as fully supervised as it simply learns DEM labels from expert-generated LBL-DEM.

#### 2.2.1. DEP Generative Adversarial Network (DEP-GAN)

DEP Generative Adversarial Network (DEP-GAN) (Rachmadi et al., 2019) is based on a GAN, a well established deep neural network model commonly used to generate fake natural images (Goodfellow et al., 2014). Thus, in this study, DEP-GAN is mainly proposed to predict the evolution of WMH when there are no longitudinal WMH labels available. DEP-GAN is based on a visual attribution GAN (VA-GAN), originally proposed to detect atrophy in T2-weighted MRI of Alzheimer’s disease (Baumgartner et al., 2017). DEP-GAN consists of a generator based on a U-Residual Network (UResNet) (Guerrero et al., 2018) and two separate convolutional networks used as discriminators (hereinafter will be referred as critics) which are based on the VA-GAN’s critics (Baumgartner et al., 2017). The schematic of DEP-GAN can be seen in Figure 2.

**Figure 2:**
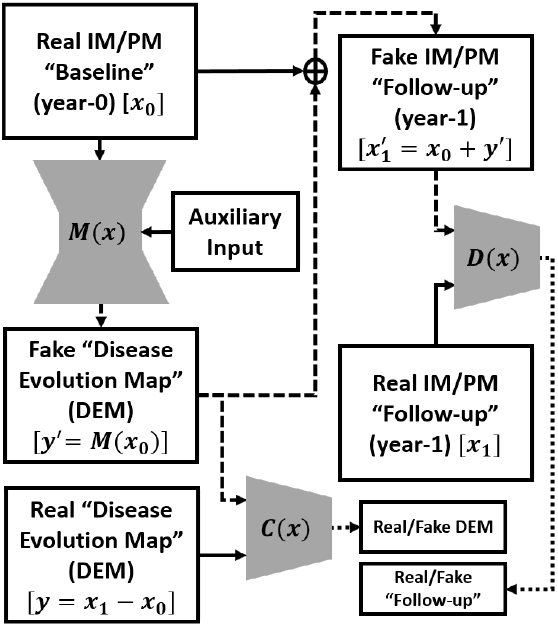
Schematic of the proposed DEP-GAN with 2 discriminators (critics). *M*(**x**) is a generator which generates “fake” disease evolution map (DEM) while *C*(**x**) and *D*(**x**) are critics to enforce anatomically realistic modifications to the follow-up images and encode realistically plausible DEMs. The flows of “fake” images are shown by the dashed lines. DEP-GAN can take either irregularity map (IM) or probability map (PM) as input. DEP-GAN also has an auxiliary input to deal with the non-deterministic factors in WMH evolution (see Section 2.3 for full explanation).

Let **x**_**0**_ be the baseline (year-0) image and **x**_**1**_ be the follow-up (year-1) image. Then, the “real” DEM (**y**) can be produced by a simple subtraction (**y** = **x**_**1**_ − **x**_**0**_). To generate the “fake” DEM (**y′**), i.e. without **x**_**1**_, a generator function (*M*(**x**)) is used: **y′** = *M*(**x**_**0**_). Thus, a “fake” follow-up image 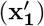 can be produced by 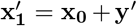. Once *M*(**x**) is well/fully trained, the “fake” follow-up 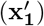 and the “real” follow-up (**x**_**1**_) should be indistinguishable by a critic function *D*(**x**), while “fake” DEM (**y′**) and “real” DEM (**y**) should be also indistinguishable by another critic function *C*(**x**). Full schematic of DEP-GAN’s architecture (i.e., its generator and critics) can be seen in Figure 3.

**Figure 3:**
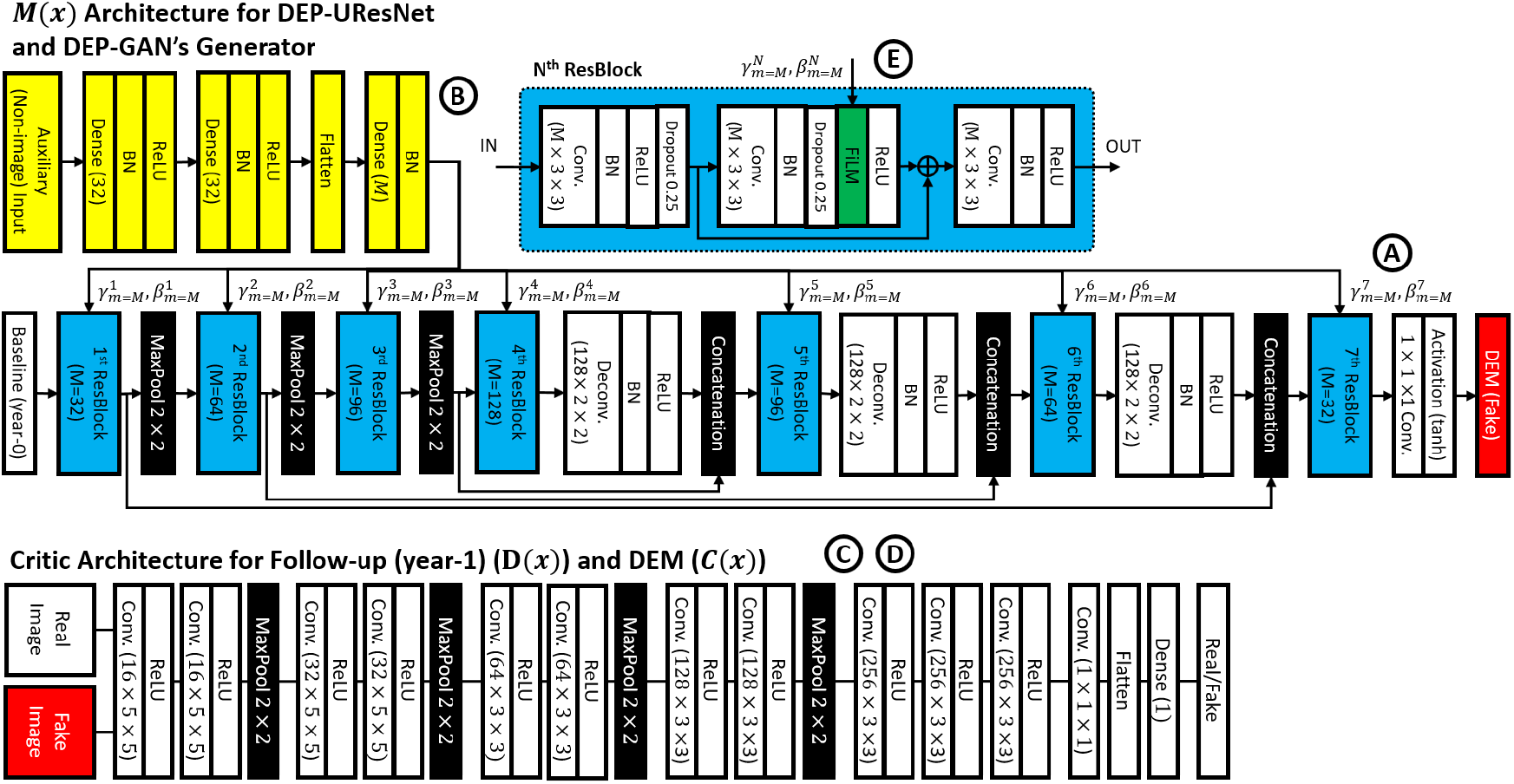
Architecture of DEP-GAN, which consists of one generator (**upper side**, “A”) and two critics (**lower side**, “C” and “D”). Note how the proposed auxiliary input is feed-forwarded to convolutional layers (yellow, “B”) and then modulated to the generator using FiLM layer (green) inside residual block (ResBlock) (light blue, “E”). Please see Figure 2 for connections between each part and Section 2.3 for full explanation about auxiliary input. On the other hand, DEP-UResNet is based on DEP-GAN’s generator, including its auxiliary input, with modification of the last non-linear activation function (i.e., use *softmax* for segmentation instead of *tanh*).

The DEP-GAN’s UResNet-based generator (*M*(**x**)) has two parts, an encoder which encodes the input image information to a latent representation and a decoder which decodes back image information from the latent representation. The baseline IM/PM (**x**_**0**_) is feed-forwarded to this generator to generate a “fake” DEM (**y′**). There is also an auxiliary input modulated into the generator using a FiLM layer (Perez et al., 2018) inside the residual block (ResBlock) to deal with non-deterministic factors of WMH evolution. This auxiliary input and its modulation will be fully discussed in Section 2.3. The architecture of the DEP-GAN’s generator is depicted in the upper side of Figure 3 (with “A”, “B”, and “E” annotations for UResNet-based generator of *M*(**x**), auxiliary input, and residual block (ResBlock) respectively).

Unlike VA-GAN that uses only one critic (i.e., only *D*(**x**)) (Baumgartner et al., 2017), DEP-GAN uses two critics (i.e., *D*(**x**) and *C*(**x**)) to enforce anatomically realistic modifications to the follow-up images (Baumgartner et al., 2017) and encode realistic plausibility in the modifier (i.e., DEM) (Rachmadi et al., 2019). Anatomically realistic modifications to the follow-up images can be achieved by optimising the critic *D*(**x**) and the anatomically realistic plausibility of the modifier can be achieved by optimising the critic *C*(**x**). In other words, we argue that an anatomically realistic DEM is essential to produce anatomically realistic (fake) follow-up images. The architecture of the DEP-GAN’s critics and their connection to the generator are depicted in the lower side of Figure 3 (with “C” and “D” annotations for critic *C*(**x**) and *D*(**x**) respectively).

The DEP-GAN’s optimisation process is the same as the optimisation of VA-GAN, where the optimisation processes of Wasserstein GAN (WGAN-GP) using a gradient penalty factor of 10 is used (Gulrajani et al., 2017). The optimisation of *M*(**x**) is given by the following function

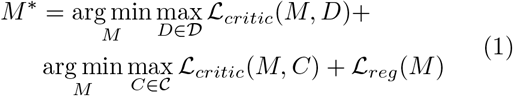

where

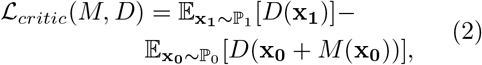

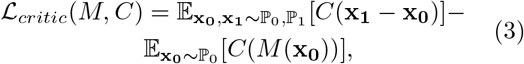

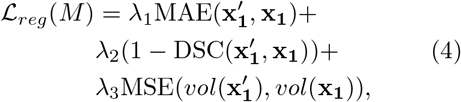

**x**_**0**_ is the baseline image that has an underlying distribution ℙ_0_, **x**_**1**_ is the follow-up image that has an underlying distribution ℙ_1_, *M*(**x**_**0**_) represents the “fake” DEM, 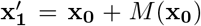 is the “fake” follow-up image, 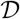 and 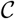 are the critics (i.e. a set of 1-Lipschitz functions (Baumgartner et al., 2017; Gulrajani et al., 2017)), and MAE and MSE are mean absolute error and mean squared error (i.e., L1 and L2 losses) respectively. The optimisation is performed by updating the parameters of the generator and critics alternately, where (each) critic is updated 5 times per generator update. Also, in the first 25 iterations and every 100 iterations, the critics are updated 100 times per generator update (Baumgartner et al., 2017; Gulrajani et al., 2017).

In summary, to optimise the generator (*M*(**x**)), we need to optimise Equation 1, which optimises both critics (*D*(**x**) and *C*(**x**)) using Equations 2 and 3 respectively based on WGAN-GP’s optimisation process (Gulrajani et al., 2017), and use the regularisation function described in Equation 4. Each term in the Equation 4 simply says:

1. Intensities of “fake” follow-up images 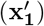 have to be similar to the “real” follow-up images (**x**_**1**_) based on MAE (i.e., L1 loss).
2. The WMH segmentation estimated from 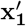 has to be spatially similar to the WMH segmentation estimated from **x**_**1**_ based on the Dice similarity coefficient (DSC) (see Equation 6).
3. The WMH volume (in *ml*) estimated from 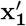 has to be similar to the WMH volume estimated from **x**_**1**_ based on MSE (i.e., L2 loss).

The WMH segmentation of 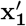 and **x**_**1**_ is estimated by either thresholding IM values (i.e., irregularity values) to be above 0.178 (Rachmadi et al., 2020) or PM values (i.e., probability values) to be above 0.5. Furthermore, each term in Equation 4 is weighted by λ_1_, λ_2_, and λ_3_ which equals to 100 (Baumgartner et al., 2017), 1, and 100 respectively.

Compared to the other GAN architectures reviewed in (Yi et al., 2019; Kazeminia et al., 2018), DEP-GAN is closely related to conditional GAN (Mirza and Osindero, 2014) with WGAN-GP’s training scheme (Gulrajani et al., 2017), both of which are the basis of the VA-GAN. The use of two discriminator/critics is also not unheard of as some relevant studies have proposed similar idea (Lanfredi et al., 2019). In regards to the application of GAN, DEP-GAN is closer to cross modality transformation than segmentation because the DEP-GAN’s generator transforms the input of IM/PM into a different modality of DEM. The uniqueness of DEP-GAN compared to other GAN architectures is the use of auxiliary input for modulating non-image information, especially the use of Gaussian noise for simulating the non-deterministic nature of WMH evolution (see Section 2.3). While modulating different modalities into deep neural networks is not entirely new, this study shows that the proposed auxiliary input improves the performance on predicting the evolution of WMH (see Section 4.2).

#### 2.2.2. DEP U-Residual Network (DEP-UResNet)

In the case of WMH binary labels (LBL) for both time points (i.e., baseline and follow-up in longitudinal data set) are available, a simple supervised deep neural network method can be used to automatically estimate WMH evolution. As previously described in Section 2.1, DEM produced from LBL (i.e., three-class DEM label (LBL-DEM)) consists of 3 foreground labels (i.e., “Grow” (green), “Shrink” (red), and “Stable” (blue)) and 1 background label (black). An example of LBL-DEM can be seen in the bottom-right figure of Figure 1.

In this study, the DEP-GAN’s generator is detached from the critics and modified into DEP U-Residual Network (DEP-UResNet) by changing the last non-linear activation layer of *tanh* (i.e., for regression) to *softmax* (i.e., for multi-label segmentation). Thus, the DEP-UResNet’s schematic is similar to the DEP-GAN’s generator, which can be seen in Figure 3 (with “A”, “B”, and “E” annotations). DEP-UResNet uses T2-FLAIR as input and LBL-DEM as target output. Note that this configuration makes all DEP models have similar generator networks based on UResNet. Furthermore, the auxiliary input proposed in this study can be also applied to the DEP-UResNet.

### 2.3. Auxiliary Input in DEP Model

The biggest challenge in modelling the evolution of WMH is mainly the amount of factors involved in WMH evolution. In our previous work, we proposed an auxiliary input module which modulates random noises from normal (Gaussian) distribution to every layer of the DEP-GAN’s generator to simulate the unknown/missing factors (i.e., non-image features) involved in WMH evolution and the non-deterministic property of WMH evolution (Rachmadi et al., 2019). To modulate the auxiliary input to every layer of the DEP-GAN’s generator we used Feature-wise Linear Modulation (FiLM) layer (Perez et al., 2018). The FiLM layer is depicted as the green block inside the residual block (ResBlock) in Figure 3 (annotated as “E”). In the FiLM layer, *γ*_*m*_ and *β*_*m*_ modulate feature maps *F*_*m*_, where subscript *m* refers to *m*^*th*^ feature map, via the following affine transformation

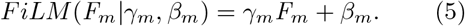

where *γ*_*m*_ and *β*_*m*_ for each ResBlock in each layer are automatically determined by convolutional layers (depicted as yellow blocks in Figure 3 with “B” annotation). Note that the proposed auxiliary input module can be easily applied to any deep neural network model. Thus, we applied the auxiliary input module to the two DEP models proposed in the present study: DEP-GAN and DEP-UResNet.

In this study, we performed an ablation study of auxiliary input modalities for DEP model by using: 1) no auxiliary input, 2) baseline WMH volume, 3) both baseline WMH and SL volumes, and 4) Gaussian noise. The WMH and SL volumes were obtained from WMH and SL labels/masks (see Section 3.1). Whereas, an array of 32 random noises which follow Gaussian distribution (Gaussian noise) of 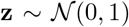 was used as per our previous work (Rachmadi et al., 2019). It is worth to mention that changing the auxiliary input modality from WMH and SL loads to Gaussian noise changes the nature of the DEP model from deterministic to non-deterministic (i.e., probabilistic).

## 3. Data and Experiments

### 3.1. Subjects and Data

We used MRI data from stroke patients (*n* = 152) enrolled in a study of stroke mechanisms from which full recruitment and assessments have been published (Wardlaw et al., 2017b). Written informed consent was obtained from all patients on protocols approved by the Lothian Ethics of Medical Research Committee (REC 09/81101/54) and NHS Lothian R+D Office (2009/W/NEU/14), on the 29th of October 2009. In the clinical study that provided the data, patients were imaged at three time points (i.e., first time (baseline) 1-4 weeks after presenting to the clinic with stroke symptoms, at approximately 3 months, and a year after (follow-up)). All images were acquired at a GE 1.5T MRI scanner following the same imaging protocol (Valdés Hernández et al., 2015). Ground truth segmentations were performed using a multi-spectral semi-automatic method (Valdés Hernández et al., 2015) only from baseline and 1-year follow-up scan visits in the image space of the T1-weighted scan of the second visit, in *n* = 152 (out of 264) patients. T2-weighted, FLAIR, gradient echo, and T1-weighted structural images at baseline and 1-year scan visits were rigidly and linearly aligned using FSL-FLIRT (Jenkinson et al., 2002). The resulted resolution of the images is 256 × 256 × 42 with slice thickness of 0.9375 × 0.9375 × 4 mm. We used data from all patients who had the three scan visits and ground truth generated as per above. Hence, our sample consists on MRI data (i.e., *s* = *n* × 2 = 304 MRI scans) for *baseline* and *1-year follow-up* data. Out of all patients, there are 70 of them that have stroke subtype lacunar (46%) with median small vessel disease (SVD) score of 1. Other demographics and clinical characteristics of the patients that provided data for this study can be seen in Table 1.

**Table 1:** Demographics and clinical characteristics of the samples used in this study (*n* = 152). SVD and PV stand for small vessel disease and periventricular respectively.

|  |  |  |
| --- | --- | --- |
| Vascular risk factors | Diabetes (n, (%)) | 18 (12) |
|  | Hypertension (n, (%)) | 114 (75) |
|  | Hypercholesterolaemia (n, (%)) | 86 (57) |
|  | Recent or present smoker (n, (%)) | 96 (64) |
| Relevant SVD imaging markers | Presence of at least 1 microbleed (n, (%)) | 26 (17) |
|  | Presence of a previous lacune (n, (%)) | 37 (24) |
|  | SVD score (median [IQR]) | 1 [0 2] |
|  | PV WMH Fazekas score (median [IQR]) | 1 [1 2] |
|  | Deep WMH Fazekas score (median [IQR]) | 1 [1 2] |

The primary study that provided the data used a semi-automatic multi-spectral method to produce several brain masks including intracranial volume (ICV), cerebrospinal fluid (CSF), stroke lesions (SL), and WMH, all which were visually checked and manually edited by an expert (Valdés Hernández et al., 2015). The image processing protocol followed to generate these masks is fully explained in (Valdés Hernández et al., 2015). Extracranial tissues, SL, and skull were removed from the baseline and follow-up T2-FLAIR images using the SL and ICV binary masks from previous analyses (Chappell et al., 2017; Wardlaw et al., 2017b). Furthermore, binary WMH labels produced for the primary study that provided the data (Valdés Hernández et al., 2015) were used as the gold standard (i.e. ground truth) for evaluating the DEP models. As per these labels, 98 and 54 out of the 152 subjects have increasing and decreasing volume of WMH respectively.

As previously explained, IM and PM are needed for DEP-GAN. We used LOTS-IM with 128 target patches (Rachmadi et al., 2020) to generate IM from each MRI data. To generate PM, we trained a 2D UResNet (Guerrero et al., 2018) with gold standard WMH and SL masks for WMH and SL segmentation. For this training, we used all subjects in our data set and a 4-fold cross validation training scheme. See Section 3.2 to see how the 4-fold cross validation is done for this study. Furthermore, note that this UResNet is different from the DEP-UResNet, which is the newly proposed model in this study. Notice that we affix “DEP” key-word to any model’s name used for prediction and delineation of WMH evolution.

### 3.2. Experiment Setup

For the present study, we opted to use 2D architectures for all our networks rather than 3D ones because the number of data available in this study is limited (i.e. only 152 subjects). VA-GAN (i.e., the GAN scheme used as basis for DEP-GAN) used roughly 4,000 subjects for training its 3D network architecture, yet there was still an evidence of over-fitting (Baumgartner et al., 2017). The 2D version of VA-GAN has been previously tested on synthetic data (Baumgartner et al., 2017).

To train DEP models (i.e., DEP-GAN and DEP-UResNet) and also UResNet (i.e., for generating PM), 4-fold cross validation was performed. Note that cross validation was not used in the previous study that introduced DEP-GAN (Rachmadi et al., 2019). In each fold, out of 304 MRI data (152 subjects × 2 scans), 228 MRI data (114 subjects × 2 scans) were used for training and 76 MRI data (38 subjects × 2 scans) were used for testing. Note that DEP models are subject-specific models, so pairwise MRI scans (i.e., baseline and follow-up) are needed and necessary for both training and testing. Out of all slices from the training set in each fold (i.e., 114 pairwise MRI scans), 20% of them were randomly selected for validation. Furthermore, we omitted slices without any brain tissues. Thus, around 4,000 slices were used in the training process in each fold. For further regularisation, we performed geometrical data augmentations (i.e., flip and rotation) and used dropout layers inside the ResBlock (see Figure 3 “E”). Values of IM/PM did not need to be normalised as these are between 0 and 1. Finally, each DEP model was trained for 200 epochs (i.e., 200 generator updates for DEP-GAN).

In this study, we first performed an ablation study using different GAN architectures for DEP model, which are based on WGAN-GP, VA-GAN, DEP-GAN with 1 critic (DEP-GAN-1C), and DEP-GAN with 2 critics (DEP-GAN-2C). This ablation study is intended to see the impact of the number of critics, the location of the critic(s), and the additional losses proposed in this study. WGAN-GP only generates DEM and has one critic for DEM (*C*(**x**)). VA-GAN and DEP-GAN-1C generate both: DEM and the follow-up image, but only have one critic for generating the follow-up image (*D*(**x**)). The difference between VA-GAN and DEP-GAN-1C is that DEP-GAN-1C has additional losses for optimisation in the training (see Section 2.2.1). Lastly, DEP-GAN-2C has two critics (*C*(**x**) and *D*(**x**)) and additional losses for the training. In this ablation study, all methods used IM and PM as main input modality and did not use any auxiliary input.

Furthermore, we also performed an ablation study using different types of auxiliary input and studied their effects to the DEP models (i.e., DEP-UResNet, DEP-GAN-2C using IM, and DEP-GAN-2C using PM). Note that we used DEP-GAN-2C for the rest of the study. The procedure of using auxiliary input depends on the input modality and training/testing process. If SL and WMH volumes were used as auxiliary input, these (i.e., not the volumes per slice, but the volume per subject) were feed-forwarded together with one MRI slice. Thus, all slices from one subject used the same number of WMH and stroke lesion volumes. Note that WMH and SL loads for the whole data set (i.e., all subjects) were first normalised to zero mean unit variance before their use in training/testing.

If Gaussian noise were used as auxiliary input, an array of Gaussian noise was feed-forwarded together with an MRI slice in the training process as follows: 10 different sets of Gaussian noise were first generated and only the “best” set (i.e., the set that yielded the lowest *M*\* loss (Equation 1)) was used to update the DEP model’s parameters. Note that this approach is similar to and inspired by Min-of-N loss in 3D object reconstruction (Fan et al., 2017) and variety loss in Social GAN (Gupta et al., 2018). In the testing process, 10 different sets of Gaussian noise were generated and the average performance was calculated. Furthermore, in the evaluation, the “best” prediction of WMH evolution based on Dice similarity coefficient (DSC) was also reported.

### 3.3. Evaluation Measurements

In this study, we used the following tests to assess the performance of DEP models:

1. **Prediction error** of WMH volumetric change (i.e., whether WMH volume in a subject will increase or decrease).
2. **Volumetric agreement** between ground truth and predicted WMH volumes of the follow-up assessment using Bland-Altman plot (Bland and Altman, 1986).
3. **Volumetric correlation** between ground truth and predicted WMH volumes of the follow-up assessment.
4. **Spatial agreement** of the automatic map of WMH evolution in a patient (i.e. after binarisation) using Dice similarity coefficient (DSC) (Dice, 1945).
5. **Clinical plausibility test** between the outcome of DEP models in relation with baseline WMH load and clinical risk factors of WMH evolution suggested in clinical studies.

**Prediction error** is a simple measurement to assess how good a DEP model can predict the WMH evolution in the future follow-up assessment (i.e., increasing or decreasing). On the other hand, **volumetric agreement** using Bland-Altman plot presents the mean volumetric difference and upper/lower limit of agreements (i.e., mean ± 1.96 × standard deviation) between ground truth and predicted WMH volumes of the follow-up assessment. We also calculated the **Volumetric correlation** between ground truth and follow-up predicted WMH volumes, complementary to the Bland-Altman plot. Whereas, for evaluating the **spatial agreement** between ground truth and automatic delineation results, we used the Dice similarity coefficient (DSC). Higher DSC means better performance, and it can be computed as follow:

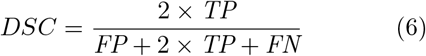

where *TP* is true positive, *FP* is false positive and *FN* is false negative.

In addition, we performed **clinical plausibility test** which evaluate the outcome of DEP models in relation with the baseline WMH load and clinical risk factors of WMH change and evolution suggested in clinical studies. For this, analyses of co-variance (ANCOVA) were performed as follows:

1. The WMH volume at follow-up, predicted from each of the schemes evaluated was used as outcome (dependent) variable.
2. The baseline WMH volume was the independent variable or predictor.
3. After running Belsley collinearity diagnostic tests, the covariates in the models were: 1) type of stroke (i.e. lacunar or cortical), 2) basal ganglia perivascular spaces (BG PVS) score, 3) presence/absence of diabetes, 4) presence/absence of hypertension, 5) recent or current smoker status (yes/no), 6) volume of the index stroke lesion (abbreviated as “index SL”), and 7) volume of old stroke lesions (abbreviated as “Old SL”).

The outcome from an ANCOVA model using the baseline and follow-up WMH volumes of the gold-standard expert-delineated binary masks was used as reference to compare the outcome of the ANCOVA models that used the volumes generated by thresholding the input and output of the DEP models. All volumetric measurements involved in the ANCOVA models were previously adjusted by patient’s head size. Therefore, all ANCOVA models used the percentage of these volumetric measurements in ICV rather than the raw volumes.

## 4. Results and Discussions

### 4.1. Ablation study of different GAN architectures for DEP model

In this ablation study, we used different GAN architectures for our DEP model to evaluate the impact of the number of critics, location of critic(s), and additional losses. See the third paragraph of Section 3.2 for full explanation of the experiments.

#### 4.1.1. Spatial agreement (DSC) and qualitative (visual) analyses

Based on Table 2 (columns 8-13), we can see that DEP-GAN-2C produced better spatial agreement (i.e., higher DSC score) than WGAN-GP, VA-GAN, and DEP-GAN-1C, especially for changing and growing WMH. Qualitative (visual) assessment of generated DEM depicted in Figure 4 also shows that DEP-GAN-2C produced more detailed DEM than the other methods, especially when compared to VA-GAN. These results show that DEP-GAN-1C and DEP-GAN-2C are more responsive to the changes of WMH and better in predicting the changes of WMH than VA-GAN. Furthermore, we also can see from both Table 2 and Figure 4 that the use of PM produced better spatial agreement than IM, regardless of the GAN architecture.

**Figure 4:**
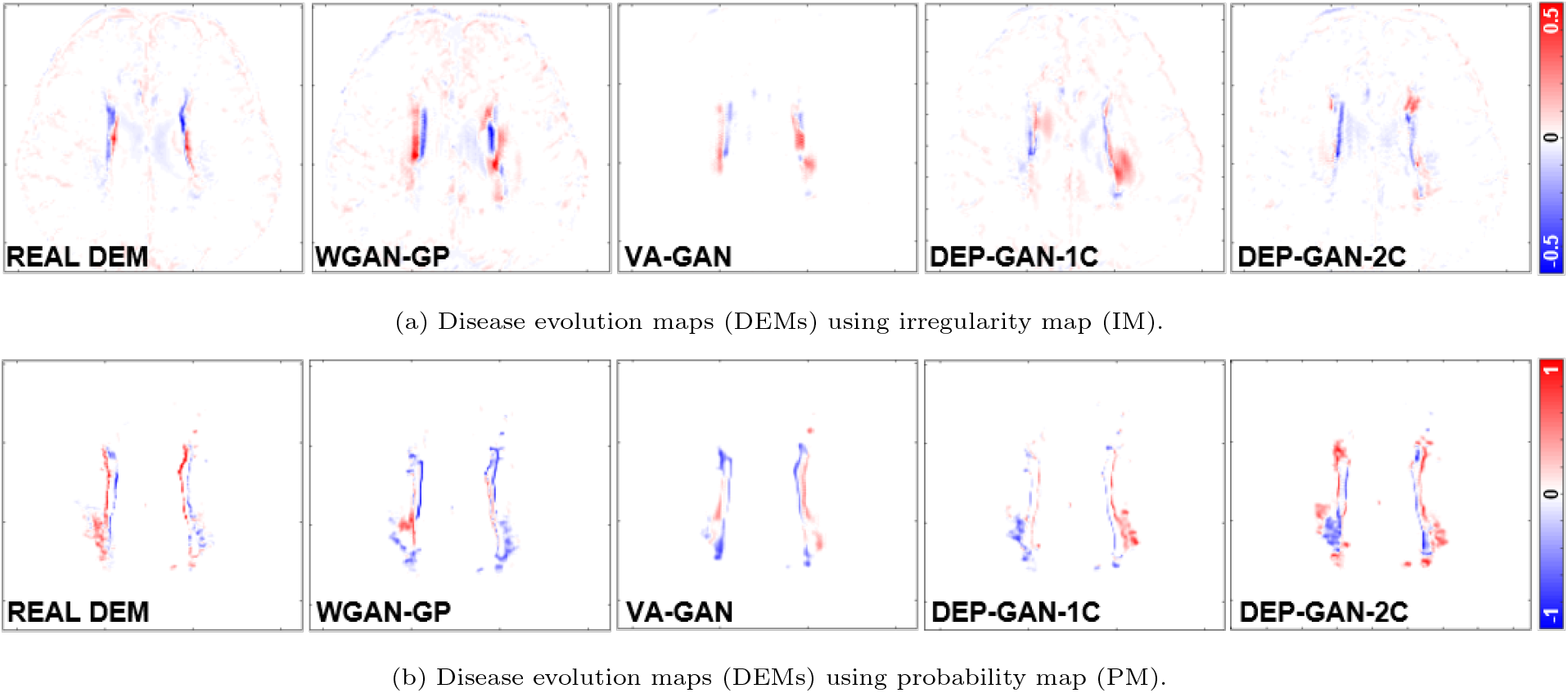
Examples of real DEM and generated DEMs produced by different GAN architectures for DEP model. From left to right: real DEM and generated DEMs produced by WGAN-GP, VA-GAN, DEP-GAN with 1 critic (DEP-GAN-1C), and DEP-GAN with 2 critics (DEP-GAN-2C) respectively.

**Table 2:** Results from ablation study of different GAN architectures for DEP models. We calculated the prediction error of WMH change, volumetric agreement of WMH volume, and spatial agreement of WMH evolution, compared to the gold standard expert-delineated WMH masks (i.e., three-class DEM labels). “DSC” stands for similarity coefficient, “Vol.” stands for volumetric, “LoA” stands for limit of agreement, and “G” and “S” stand for percentage of subjects correctly predicted as having growing and shrinking WMH by DEP models. The best value for each learning approaches and evaluation measurements is written in bold.

| Irregularity Map | Grow (G) [%] | Shrink (S) [%] | Avg. [%]<br>((G+S)/2) | Vol. Bias [ml]<br>mean(std) | Lower LoA [ml] | Upper LoA [ml] | Entire WMH | Change (C) | Stable (St) | Shrink (Sr) | Grow (Gr) | Avg. ((St+Sr+Gr)/3) |
| --- | --- | --- | --- | --- | --- | --- | --- | --- | --- | --- | --- | --- |
| WGAN-GP | **85.71** | 40.74 | 63.23 | -11.70(24.12) | -59.11 | 35.70 | 0.3179 | 0.0809 | 0.3294 | **0.0595** | 0.0325 | 0.1405 |
| VA-GAN | 65.31 | 62.96 | 64.13 | **2.52(16.43)** | -29.69 | **34.72** | **0.3361** | 0.0789 | 0.3506 | 0.0356 | 0.0361 | 0.1408 |
| DEP-GAN-1C | 65.31 | 68.52 | **66.91** | 3.88(15.93) | -27.33 | 35.10 | 0.3343 | 0.0583 | **0.3711** | 0.0388 | 0.0265 | 0.1454 |
| DEP-GAN-2C | 61.22 | **72.22** | 66.72 | 5.54(15.98) | **-25.79** | 36.87 | 0.3204 | **0.0946** | 0.3684 | 0.0238 | **0.0445** | **0.1456** |
| Probability Map | Grow (G) [%] | Shrink (S) [%] | Avg. [%]<br>((G+S)/2) | Vol. Bias [ml]<br>mean(std) | Lower LoA [ml] | Upper LoA [ml] | Entire WMH | Change (C) | Stable (St) | Shrink (Sr) | Grow (Gr) | Avg. ((St+Sr+Gr)/3) |
| WGAN-GP | 55.10 | 79.63 | 67.37 | 4.19(8.28) | -12.05 | 20.42 | **0.6139** | 0.2082 | 0.5906 | 0.1494 | 0.0899 | 0.2766 |
| VA-GAN | 42.86 | **94.44** | 68.65 | 5.78(8.13) | **-10.15** | 21.70 | 0.6070 | 0.1946 | 0.5952 | **0.1584** | 0.0641 | 0.2726 |
| DEP-GAN-1C | 59.18 | 85.19 | 72.18 | 3.66(7.64) | -11.32 | **18.63** | 0.6116 | 0.1711 | **0.6012** | 0.1186 | 0.0800 | 0.2666 |
| DEP-GAN-2C | **69.30** | 75.93 | **72.66** | **2.48(8.47)** | -14.13 | 19.08 | 0.6083 | **0.2246** | 0.5812 | 0.1515 | **0.1105** | **0.2811** |

#### 4.1.2. Volumetric agreement (Bland-Altman) and correlation analyses

From Table 3, we can see that the volume of WMH predicted by DEP-GAN-1C and DEP-GAN-2C correlated better with the volume of the ground truth than the volume of WMH predicted using WGAN-GP and VA-GAN. However, as per the volumetric agreement (Bland-Altman) analysis, the performance of DEP-GAN-1C and DEP-GAN-2C depended on the working domain, IM or PM (see columns 5-7 of Table 2). If PM was used, DEP-GAN-1C and DEP-GAN-2C performed better than the other methods. On the other hand, VA-GAN achieved the best volumetric agreement when IM was used. However, VA-GAN’s good performance in the volumetric agreement analysis did not translate to good spatial agreement as previously described in Section 4.1.1.

**Table 3:** Volumetric correlation analysis in ablation study of GAN architectures for DEP model. The best value for each correlation measurement is written in bold.

| Irregularity Map | WGAN-GP | VA-GAN | DEP-GAN-1C | DEP-GAN-2C |
| --- | --- | --- | --- | --- |
| R<sup>2</sup> | 0.1394 | 0.5644 | 0.5999 | **0.6068** |
| Trend | $y = 0.3354x + 6.5866$ | $y = 0.4056x + 2.7858$ | $y = \mathbf{0.4225x} + \mathbf{2.3714}$ | $y = 0.4159x + 2.0128$ |
| Probability Map | WGAN-GP | VA-GAN | DEP-GAN-1C | DEP-GAN-2C |
| R<sup>2</sup> | 0.8735 | 0.8813 | **0.8916** | 0.8659 |
| Trend | $y = 0.8525x - 0.1265$ | $y = 0.8289x - 0.3792$ | $y = 0.8799x - 0.1667$ | $y = \mathbf{0.898x} + \mathbf{0.0258}$ |

Based on the Bland-Altman and correlation plots depicted in Figure 5, we can see that PM is better than IM for representing the volumetric change of WMH where the correlation between ground truth and predicted WMH volumes when PM was used is higher than when IM was used, regardless of the GAN architecture. Furthermore, Bland-Altman plots show evidence of increasing discrepancy and variability between ground truth and predicted volumes with increasing volume of WMH when IM was used. These discrepancy and variability are less prominent when PM was used.

**Figure 5:**
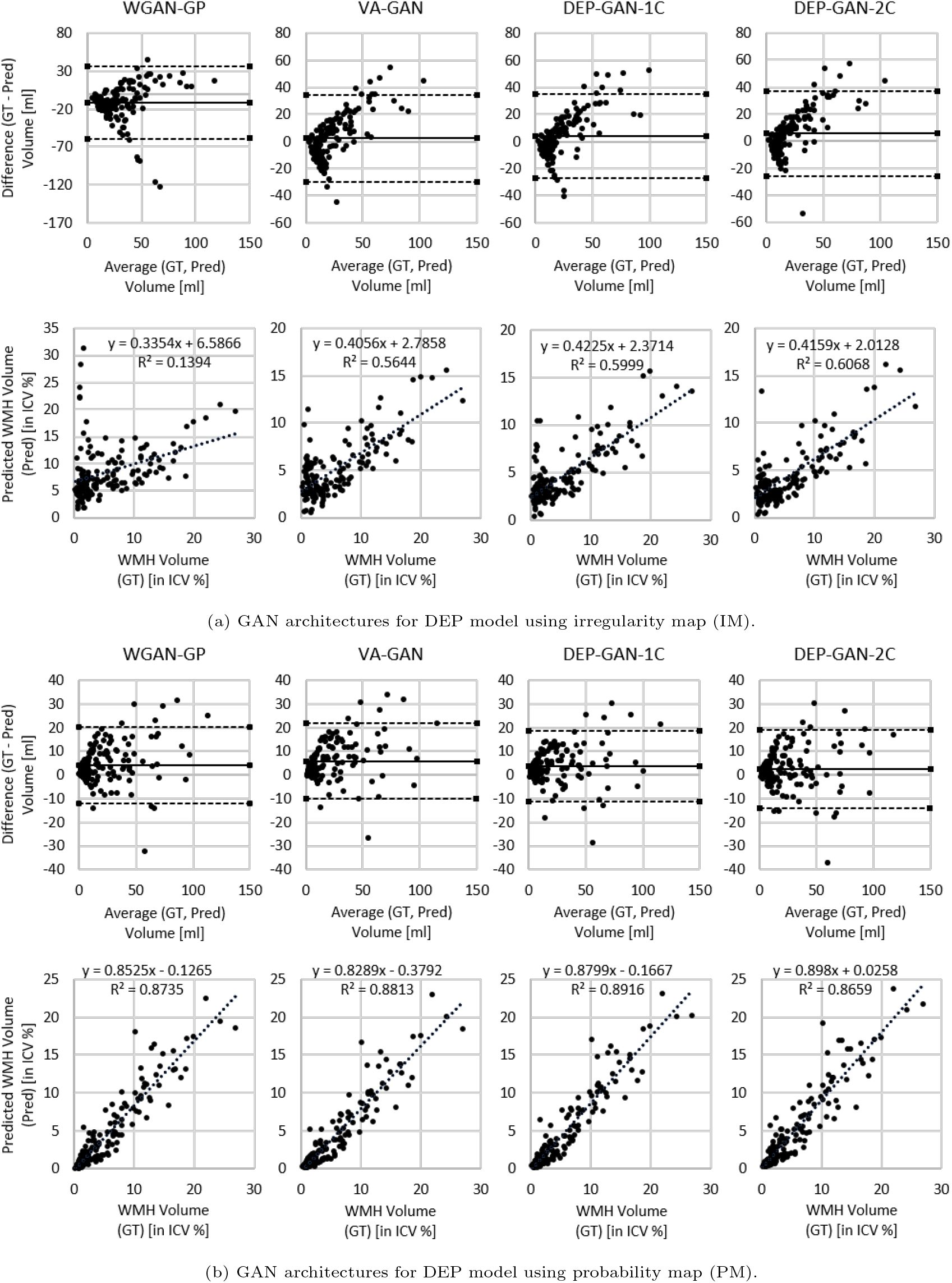
Volumetric agreement (in *ml*) and correlation (in ICV %) analyses between ground truth (GT) and predicted volume of WMH (Pred) produced by WGAN-GP, VA-GAN, DEP-GAN-1C, and DEP-GAN-2C using (a) IM and (b) PM using Bland-Altman and correlation plots.

#### 4.1.3. Prediction error analysis and discussion

From Table 2 (columns 2-4), we can see that most GAN-based DEP models could correctly predict the progression/regression of WMH volume, as they performed better than a random guess system (≥ 50%). Furthermore, we can conclude that DEP-GAN-2C performed generally better for predicting the evolution of WMH due to additional losses and two critics in the architecture. There is also evidence that PM is better for representing the evolution of WMH than IM when GAN-based deep learning methods are used.

### 4.2. Ablation study of auxiliary input in DEP models

In this ablation study, we used different types (modalities) of auxiliary input to see how they affect the performance of DEP models for predicting the evolution of WMH. Please read the fourth and fifth paragraphs of Section 3.2 for full explanation of the experiments.

#### 4.2.1. Volumetric agreement (Bland-Altman) and correlation analyses

From Table 4 (columns 5-7), DEP-UResNet using Gaussian noise (+Gaussian (mean)) produced the best estimation of WMH volumetric changes. Also, almost all DEP-UResNet models with auxiliary input performed better in volumetric agreement analysis than ones without auxiliary input. Only DEP-UResNet with WMH performed slightly lower than DEP-UResNet without auxiliary input. This shows the importance of auxiliary input for predicting the evolution of WMH using deep neural networks.

**Table 4:** Results from ablation study of auxiliary input in DEP models. Prediction error of WMH change, volumetric agreement of WMH volume, and spatial agreement of WMH evolution were calculated to the gold standard expert-delineated WMH masks (i.e., three-class DEM labels). “DSC” stands for similarity coefficient, “Vol.” stands for volumetric, “LoA” stands for limit of agreement, and “G” and “S” stand for percentage of subjects correctly predicted as having growing and shrinking WMH by DEP models. The best value for each machine learning approaches and evaluation measurements is written in bold. Furthermore, the best value of all learning approaches for each evaluation measurements is underlined and written in bold.

| DEP-UResNet | Grow<br>(G) [%] | Shrink<br>(S) [%] | Avg. [%]<br>((G+S)/2) | Vol. Bias [ml]<br>mean(std) | Lower<br>LoA [ml] | Upper<br>LoA [ml] | Entire<br>WMH | Change<br>(C) | Stable<br>(St) | Shrink<br>(Sr) | Grow<br>(Gr) | Avg. ((Sr+Gr+St)/3) |
| --- | --- | --- | --- | --- | --- | --- | --- | --- | --- | --- | --- | --- |
| No Auxiliary | 70.41 | 72.22 | 71.32 | 1.16(7.31) | **-13.17** | 15.48 | 0.6091 | 0.2234 | **0.6332** | 0.1551 | 0.1128 | 0.3004 |
| +WMH | 73.47 | **77.78** | 75.62 | 1.59(7.85) | -13.80 | 16.97 | 0.6005 | 0.2532 | 0.6188 | 0.1688 | 0.1409 | 0.3095 |
| +WMH+Stroke | 79.59 | 75.93 | **77.76** | 0.81(8.14) | -15.14 | 16.76 | 0.6080 | 0.2565 | 0.6311 | 0.1688 | 0.1415 | 0.3138 |
| +Gaussian (mean) | **81.63** | 59.26 | 70.45 | **-0.58(7.99)** | -16.24 | 15.09 | **0.6135** | **0.2629** | 0.6230 | **0.1717** | **0.1477** | **0.3141** |
| +Gaussian (best) | 81.63 | 57.41 | 69.52 | -0.79(7.96) | -16.40 | 14.81 | 0.6162 | 0.2686 | 0.6280 | 0.1787 | 0.1409 | 0.3159 |
| DEP-GAN-2C & IM | Grow<br>(G) [%] | Shrink<br>(S) [%] | Avg. [%]<br>((G+S)/2) | Vol. Bias [ml]<br>mean(std) | Lower<br>LoA [ml] | Upper<br>LoA [ml] | Entire<br>WMH | Change<br>(C) | Stable<br>(St) | Shrink<br>(Sr) | Grow<br>(Gr) | Avg. ((Sr+Gr+St)/3) |
| No Auxiliary | 61.22 | **72.22** | 66.72 | 5.58(15.98) | -25.79 | 36.87 | 0.3204 | 0.0946 | 0.3684 | 0.0238 | 0.0445 | 0.1456 |
| +WMH | **75.51** | 53.70 | 64.61 | -1.18(19.71) | -39.80 | 37.45 | 0.3249 | 0.0901 | 0.3551 | 0.0580 | 0.0458 | 0.1530 |
| +WMH+Stroke | 71.43 | 64.81 | **68.12** | **0.92(19.91)** | -38.11 | 39.95 | 0.3291 | 0.0922 | 0.3476 | **0.0590** | **0.0468** | 0.1511 |
| +Gaussian (mean) | 61.22 | 70.37 | 65.80 | 4.59(14.99) | **-24.79** | **33.98** | **0.3359** | **0.2252** | **0.3768** | 0.0485 | 0.0361 | **0.1538** |
| +Gaussian (best) | 72.45 | 64.81 | 68.83 | 0.44(15.37) | -29.67 | 30.56 | 0.3429 | 0.1053 | 0.3795 | 0.0619 | 0.0633 | 0.1682 |
| DEP-GAN-2C & PM | Grow<br>(G) [%] | Shrink<br>(S) [%] | Avg. [%]<br>((G+S)/2) | Vol. Bias [ml]<br>mean(std) | Lower<br>LoA [ml] | Upper<br>LoA [ml] | Entire<br>WMH | Change<br>(C) | Stable<br>(St) | Shrink<br>(Sr) | Grow<br>(Gr) | Avg. ((Sr+Gr+St)/3) |
| No Auxiliary | **69.39** | 75.93 | 72.66 | 2.48(8.47) | -14.13 | 19.08 | 0.6083 | 0.2246 | 0.5812 | 0.1515 | 0.1105 | 0.2811 |
| +WMH | 68.37 | 70.37 | 69.37 | **1.70(8.24)** | -14.45 | **17.84** | **0.6125** | **0.2295** | 0.6006 | 0.1467 | **0.1267** | **0.2913** |
| +WMH+Stroke | 66.33 | 75.93 | 71.13 | 2.69(9.14) | -15.22 | 20.60 | 0.6098 | 0.2229 | 0.5943 | **0.1581** | 0.1091 | 0.2872 |
| +Gaussian (mean) | 58.16 | **79.63** | **68.90** | 2.91(8.81) | **-14.36** | 20.18 | 0.6107 | 0.1801 | **0.6245** | 0.1216 | 0.0868 | 0.2776 |
| +Gaussian (best) | 65.31 | 88.89 | 77.10 | 3.63(7.85) | -11.75 | 19.02 | 0.6155 | 0.2415 | 0.6044 | 0.1834 | 0.1265 | 0.3048 |

On the other hand, from all DEP models, DEP-GAN-2C using IM produced the worst standard deviation (std) and (lower and upper) limits of agreement (LoA) in the volumetric agreement analysis, regardless of the modalities of auxiliary input. This is another indication that IM is not adequate for predicting the evolution of WMH. Interestingly, DEP-GAN-2C using PM, which seemingly had better (lower and upper) LoA than the DEP-GAN-2C using IM, had some of the worst mean of volumetric bias. This indicates that there is a bias towards regression (i.e., shrinking of WMH) when DEP-GAN-2C using PM was used for predicting the evolution of WMH. Furthermore, the correlations between ground truth and predicted volumes of WMH for DEP-UResNet and DEP-GAN-2C using PM were much higher than the ones produced by DEP-GAN-2C using IM, especially when auxiliary input is incorporated (see Table 5). All Bland-Altman and correlation plots can be found in the supplementary materials.

**Table 5:** Volumetric correlation analysis of DEP models with different types/modalities of auxiliary input in ablation study of auxiliary input.

| DEP-UResNet | R<sup>2</sup> | Trend |
| --- | --- | --- |
| No Auxiliary | **0.9031** | $y = 0.9781x - 0.1397$ |
| +WMH | 0.8893 | $y = \mathbf{1.0113x} - \mathbf{0.2435}$ |
| +WMH+Stroke | 0.8939 | $y = 0.984x - 0.2768$ |
| +Gaussian (mean) | 0.8855 | $y = 0.9772x + 0.2841$ |
| +Gaussian (best) | 0.8869 | $y = 0.9821x + 0.3073$ |
| DEP-GAN-2C & IM | R<sup>2</sup> | Trend |
| No Auxiliary | 0.6068 | $y = 0.4159x + 2.0128$ |
| +WMH | 0.3293 | $y = 0.3539x + 3.9732$ |
| +WMH+Stroke | 0.3129 | $y = 0.3817x + 3.275$ |
| +Gaussian (mean) | **0.6461** | $y = \mathbf{0.4684x} + \mathbf{1.9418}$ |
| +Gaussian (best) | 0.6037 | $y = 0.4724x + 2.9103$ |
| DEP-GAN-2C & PM | R<sup>2</sup> | Trend |
| No Auxiliary | 0.8659 | $y = 0.898x + 0.0258$ |
| +WMH | 0.8755 | $y = \mathbf{0.9541x} - \mathbf{0.1169}$ |
| +WMH+Stroke | **0.8916** | $y = 0.9102x - 0.0987$ |
| +Gaussian (mean) | 0.8541 | $y = 0.9228x - 0.23$ |
| +Gaussian (best) | 0.8836 | $y = 0.8972x - 0.2629$ |

#### 4.2.2. Spatial agreement (DSC) analysis

On the automatic delineation of WMH change’s boundaries in the follow-up year, DEP-UResNet using Gaussian noise produced the best performances for the entire WMH and the average of stable, shrinking, and growing WMH clusters (in Table 4 columns 8-13). Furthermore, it also outperformed the rest of the models on changing, shrinking, and growing WMH clusters. Compared to the “vanilla” DEP-UResNet with No Auxiliary, paired two-sided Wilcoxon signed rank tests yielded *p*-values of 0.1563, 0.0425, 0.0625, 0.0313, 0.0313, and 0.0425 for the entire WMH, changing WMH, stable WMH, shrinking WMH, growing WMH, and average respectively. These results clearly show the advantage of performing fully human supervised learning and modulating Gaussian noise as auxiliary input for predicting the evolution of WMH. It is also worth mentioning that its performance is even better when the “best” Gaussian noise is used for evaluation.

From the results in Table 4, DEP-GAN-2C using PM had close performance to the DEP-UResNet in all performed analyses, especially in the spatial agreement analysis (columns 8-13). To give a better visualisation of the spread of the performances, we plotted the distributions of DSC scores for all WMH categories using box-plots (Figure 6). Performances of DEP-GAN-2C using PM and DEP-UResNet on delineating different WMH clusters were similar in the distribution of DSC scores. Based on paired two-sided Wilcoxon signed rank tests, there was no significant difference between the performances of DEP-GAN-2C using PM and DEP-UResNet in all WMH clusters, especially when the same auxiliary input was used, with p-value > 0.17. In contrast, the differences of DSC scores in all WMH clusters produced by DEP-GAN-2C using IM and DEP-UResNet were significantly different from each other with p-value < 0.0012.

**Figure 6:**
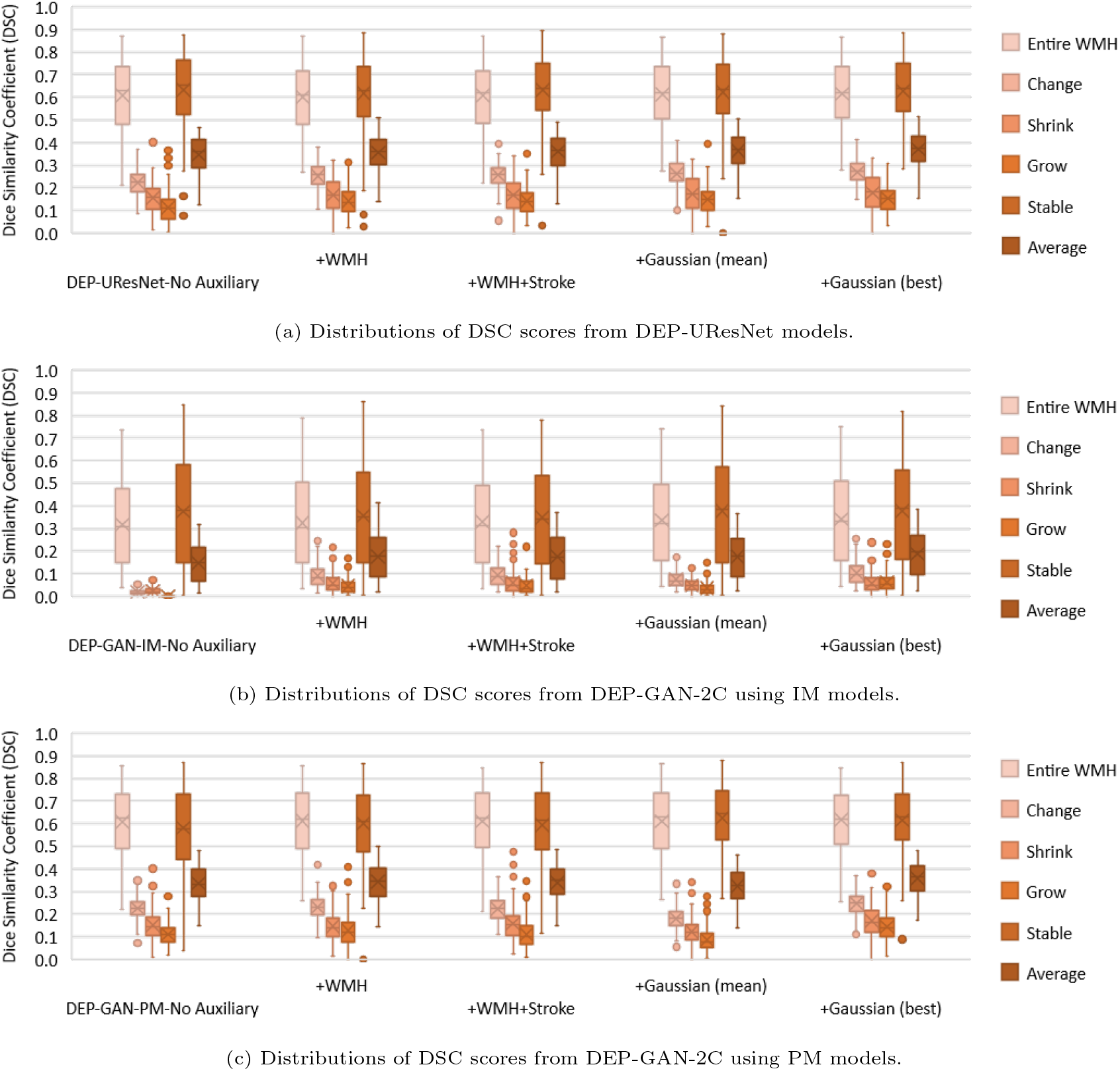
Distributions of DSC scores from all evaluated DEP models in auxiliary input ablation study. These distributions correspond to the Table 4, columns 8-13.

#### 4.2.3. Qualitative (visual) analysis

It is worth to mention first that the growing and shrinking regions of WMH are considerably smaller than those unchanged (stable) as depicted in Figure 8. Furthermore, it is very difficult to discern the borders between growing and shrinking regions when stroke lesions coalesce with WMH despite stroke lesions being removed from the analysis as previously explained. Nevertheless, inaccuracies while determining the borders between coalescent WMH and stroke lesions and the small size of the volume changes in each WMH cluster (Rachmadi et al., 2018a) might have influenced in the low DSC values obtained in the regions that experienced change as seen in Table 4. Furthermore, it is also worth to note that most regions of WMH are stable, and DEP-UResNet and DEP-GAN-2C using PM did not have any problem on segmenting these regions as depicted in Figures 8 and 9.

Based on the qualitative (visual) assessment of the DEM produced by DEP-GAN-2C using IM/PM depicted in Figure 7, auxiliary input improved the quality of the generated DEMs where they had more correct details than the ones generated without using auxiliary input. However, good details of the generated DEM from IM/PM did not necessarily translate to good three-class DEM label (i.e., three labels of growing, shrinking, and stable WMH) as depicted in Figure 9. Some reasons that might have caused this are; 1) the generated DEM from IM/PM is result of a regression process from the baseline IM/PM using DEP-GAN and 2) the three-class DEM label itself is generated from the resulted regression, where WMH is defined by having irregularity/probability values greater than or equal to 0.178 for IM and 0.5 for PM (Rachmadi et al., 2020). Note that regression of the whole brain using IM/PM is harder than direct segmentation of three regions of WMH (i.e., stable, shrinking, and growing WMH). Furthermore, small changes in IM/PM did not necessarily change the state of voxel from WMH to non-WMH or vice versa. These are the challenges of performing prediction of WMH evolution using DEP-GAN-2 and IM/PM instead of DEP-UResNet.

**Figure 7:**
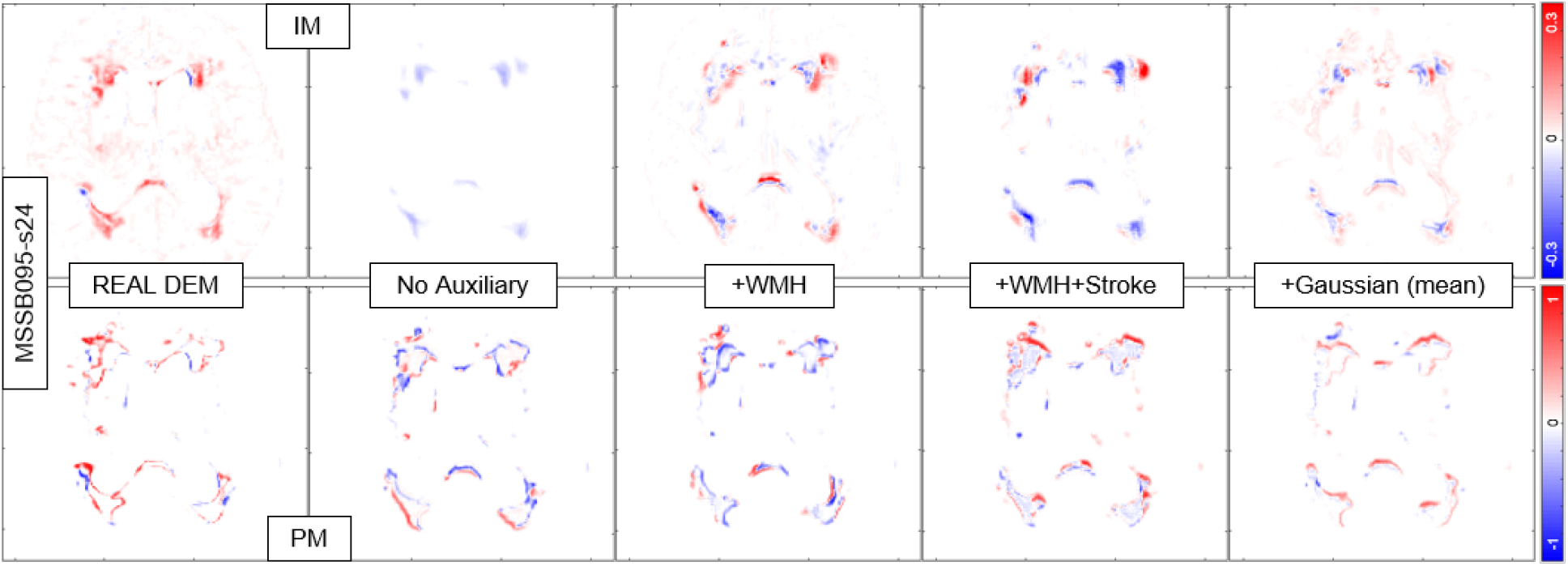
Qualitative (visual) assessment of DEM produced by DEP-GAN-2C using irregularity map (IM) and DEP-GAN-2C using probability map (PM) with different types/modalities of auxiliary input. The corresponding T2-FLAIR (input data) can be seen in Figure 9.

**Figure 8:**
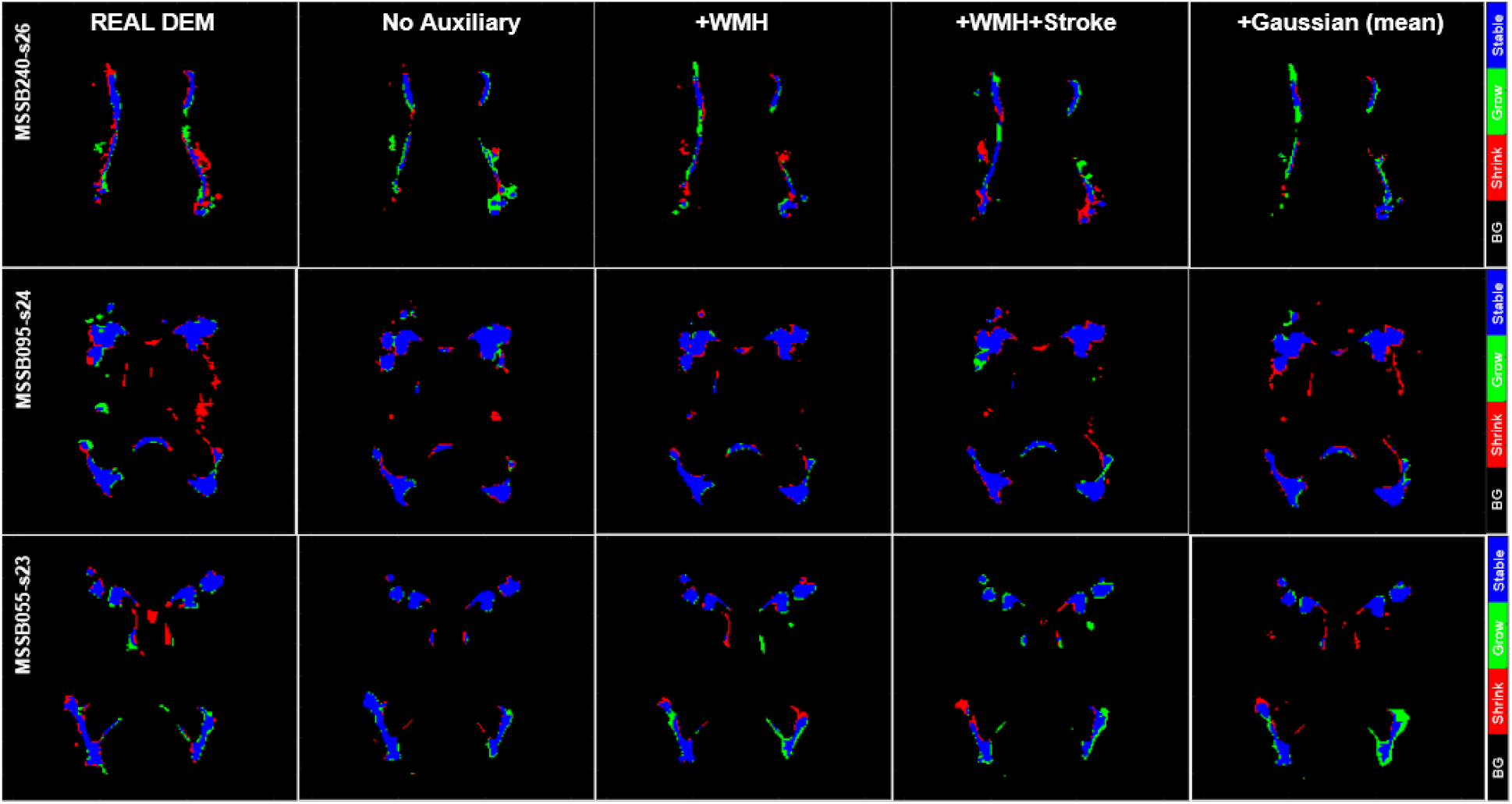
Qualitative (visual) assessment of DEM label produced by DEP-UResNet with different types/modalities of auxiliary input. The corresponding T2-FLAIR (input data) can be seen in Figure 9.

**Figure 9:**
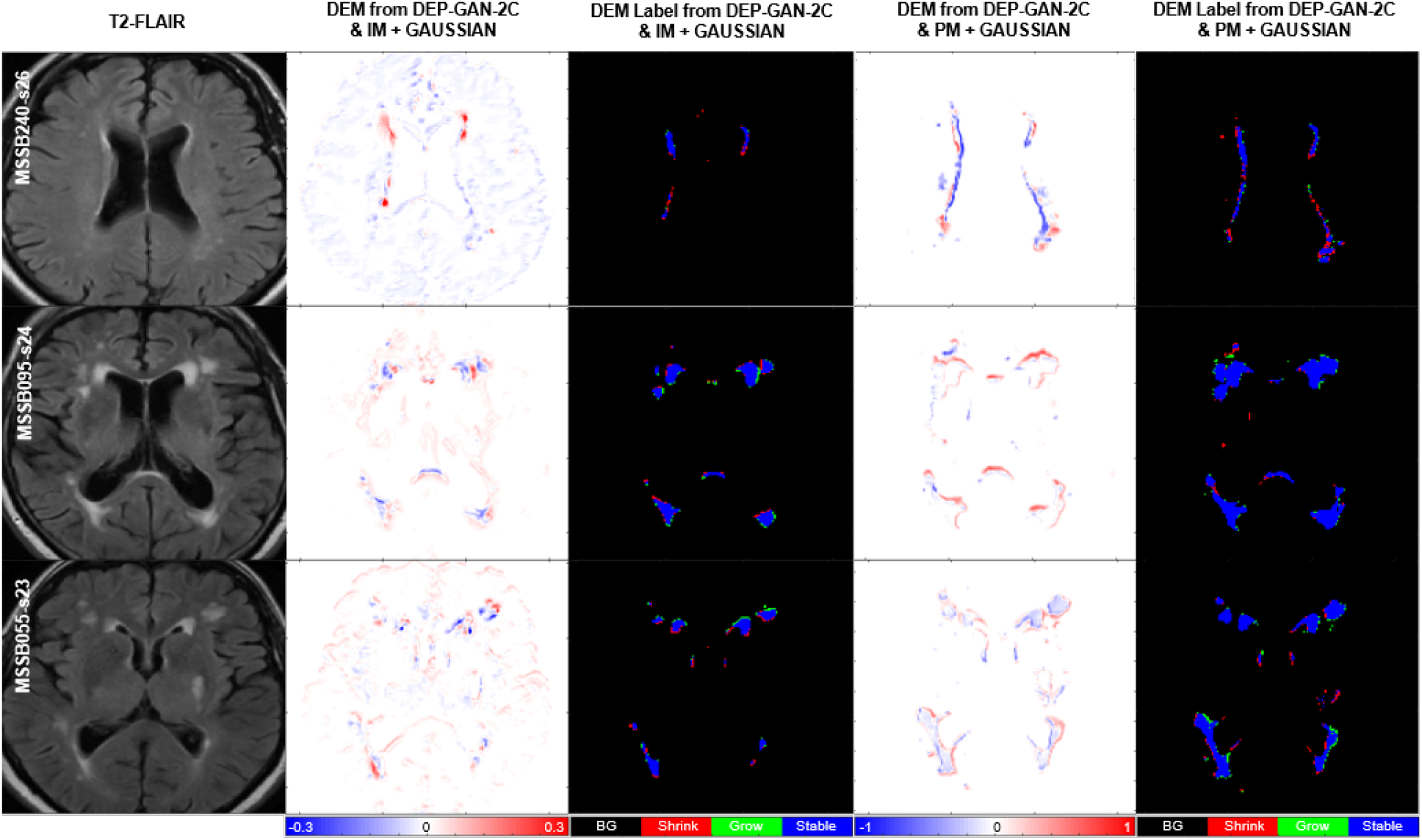
Qualitative (visual) assessment of DEM and its corresponding DEM label produced by DEP-GAN-2C using irregularity map (IM) and DEP-GAN-2c using probability map (PM) respectively, with different types/modalities of auxiliary input. The corresponding golden standard of DEM label can be seen in Figure 8.

#### 4.2.4. Clinical plausibility analysis

From Table 6, we can see that the use of expert-delineated binary WMH masks and WMH maps obtained from thresholding IM or PM (see from second to fourth rows), all produced the same ANCOVA model’s results; none of the covariates of the model had an effect in the 1-year WMH volume change, yielding almost identical numerical results in the first two decimal places. Therefore, the use of LOTS-IM and UResNet, generators of the IM and PM respectively, for producing WMH maps in clinical studies of mild to moderate stroke seems plausible.

**Table 6:** Results from the ANCOVA models that investigate the effect of several clinical variables (i.e. stroke subtype, stroke-related imaging markers and vascular risk factors) in the WMH volume change from baseline to one year after. The first column at the left hand side refers to the models/methods used to obtain the follow-up WMH volume used in the ANCOVA models as outcome variable. The rest of the columns show the coefficient estimates B and the significance level given by the p-value (i.e. B(p)), for each covariate included in the models.

| Reference<br>(binary mask) | Stroke<br>lacunar | BG PVS<br>scores | Diabetes<br>(y/n) | Hypertension<br>(y/n) | Smoker<br>(y/n) | Index SL<br>(% in ICV) | Old SL<br>(% in ICV) |
| --- | --- | --- | --- | --- | --- | --- | --- |
| Expert-delineated | -0.04(0.65) | 0.07(0.25) | -0.10(0.48) | -0.05(0.66) | -0.07(0.42) | -0.03(0.46) | 0.13(0.15) |
| Thresholded IM | -0.04(0.66) | 0.08(0.19) | -0.12(0.44) | -0.04(0.71) | -0.09(0.38) | -0.03(0.43) | 0.14(0.14) |
| Thresholded PM | -0.04(0.66) | 0.08(0.19) | -0.12(0.44) | -0.04(0.71) | -0.09(0.38) | -0.03(0.43) | 0.14(0.14) |
| DEP-UResNet | Stroke<br>lacunar | BG PVS<br>scores | Diabetes<br>(y/n) | Hypertension<br>(y/n) | Smoker<br>(y/n) | Index SL<br>(% in ICV) | Old SL<br>(% in ICV) |
| No Auxiliary | -0.12(0.11) | 0.10(0.03) | -0.06(0.57) | 0.03(0.73) | -0.08(0.29) | -0.04(0.14) | 0.30(<0.001) |
| +WMH | -0.10(0.13) | 0.11(0.006) | 0.04(0.65) | 0.01(0.87) | -0.05(0.38) | -0.04(0.13) | 0.20(<0.001) |
| +WMH+Stroke | -0.07(0.29) | 0.06(0.14) | 0.07(0.48) | -0.02(0.75) | -0.10(0.15) | -0.05(0.10) | 0.32(<0.001) |
| +Gaussian (mean) | -0.09(0.26) | 0.11(0.04) | 0.06(0.61) | 0.02(0.81) | -0.10(0.21) | -0.06(0.08) | 0.36(<0.001) |
| DEP-GAN-2C & IM | Stroke<br>lacunar | BG PVS<br>scores | Diabetes<br>(y/n) | Hypertension<br>(y/n) | Smoker<br>(y/n) | Index SL<br>(% in ICV) | Old SL<br>(% in ICV) |
| No Auxiliary | 0.03(0.68) | -0.03(0.58) | -0.07(0.54) | 0.0006(0.99) | -0.08(0.33) | -0.11(0.001) | 0.25(0.001) |
| +WMH | 0.22(0.09) | 0.08(0.36) | -0.004(0.98) | 0.12(0.40) | -0.08(0.54) | -0.06(0.25) | 0.32(0.01) |
| +WMH+Stroke | -0.11(0.45) | -0.08(0.40) | 0.03(0.88) | 0.10(0.53) | 0.11(0.47) | -0.02(0.77) | 0.34(0.02) |
| +Gaussian (mean) | -0.02(0.86) | -0.07(0.24) | -0.06(0.69) | -0.05(0.62) | -0.07(0.43) | -0.14(0.0004) | 0.20(0.03) |
| DEP-GAN-2C & PM | Stroke<br>lacunar | BG PVS<br>scores | Diabetes<br>(y/n) | Hypertension<br>(y/n) | Smoker<br>(y/n) | Index SL<br>(% in ICV) | Old SL<br>(% in ICV) |
| No Auxiliary | -0.10(0.24) | 0.14(0.009) | 0.10(0.45) | 0.04(0.67) | -0.03(0.70) | -0.05(0.18) | 0.18(0.03) |
| +WMH | -0.03(0.72) | 0.09(0.09) | -0.14(0.31) | -0.04(0.68) | -0.06(0.46) | -0.04(0.30) | 0.19(0.03) |
| +WMH+Stroke | -0.10(0.28) | 0.17(0.006) | 0.10(0.50) | 0.10(0.36) | -0.02(0.81) | -0.08(0.05) | 0.24(0.01) |
| +Gaussian (mean) | -0.09(0.25) | 0.10(0.04) | 0.02(0.87) | -0.0001(0.99) | -0.08(0.27) | -0.04(0.17) | 0.14(0.05) |

As discussed in Section 1, baseline WMH volume has been recognised the main predictor of WMH change over time (Chappell et al., 2017; Wardlaw et al., 2017b), although the existence of previous stroke lesions (SL) and hypertension have been acknowledged as contributed factors. However, from the results of the ANCOVA models (Table 6), none of the DEP models that used these (i.e WMH and/or SL volumes) as auxiliary inputs showed similar performance (i.e. in terms of strength and significance in the effect of all the covariates in the WMH change) as the reference WMH maps. The only DEP model that shows promise in reflecting the effect of the clinical factors selected as covariates in WMH progression was the DEP-GAN-2C that used as input the PM of baseline WMH and Gaussian noise (i.e. written in bold and underlined in the left hand side column of Table 6).

Some factors might have adversely influenced the performance of these predictive models. First, all deep-learning schemes require a very large amount of balanced (e.g. in terms of the appearance, frequency and location of the feature of interest, i.e. WMH in this case) data, generally not available. The lack of data available imposed the use of 2D model configurations, which generated unbalance in the training: for example, not all axial slices have the same probability of WMH occurrence, also WMH are known to be less frequent in temporal lobes and temporal poles are a common site of artefacts affecting the IM and PM, error that might propagate or even be accentuated when these modalities are used as inputs. Second, the combination of hypertension, age and the extent, type, lapse of time since occurrence and location of the stroke might be influential on the WMH evolution, therefore rather than a single value, the incorporation of a model that combines these factors would be beneficial. However, such model is still to be developed also due to lack of data available. Third, the tissue properties have not been considered. A model to reflect the brain tissue properties in combination with vascular and inflammatory risk factors is still to be developed. Lastly, the deep-learning models as we know them, although promising, are reproductive, not creative. The development of more advanced inference systems is paramount before these schemes can be used in clinical practice.

#### 4.2.5. Prediction error analysis and discussion

From Table 4 (columns 2-4), we can see that all DEP models tested in this ablation study could correctly predict the progression/regression of WMH volume better than a random guess system (≥ 50%). Furthermore, DEP models with auxiliary input, either Gaussian noise or known risk factors of WMH evolution (i.e., WMH and SL loads), produced better performances in most cases and evaluation analyses than the DEP models without any auxiliary input. These results show the importance of auxiliary input, especially Gaussian noise which simulates the non-deterministic nature of WMH evolution.

Based on our careful examinations, the Gaussian noise can change the predictions on individual subjects, but not drastically in every subject. In fact, the most probable prediction result of DEP models can be determined by the help of Gaussian noise auxiliary input. For example, subjects with high numbers of progression/regression prediction (i.e., predicted as such in more than 6 tests out of 10) can be considered of having higher probability of WMH progression/regression respectively. It is also worth mentioning that any outlier in prediction can be generalised by averaging all possible outputs for the final prediction. Thus, we can sample more than 10 times in the testing to get a better final prediction result. This generalisation cannot be done without using Gaussian noise as auxiliary input.

Furthermore, it is clear now that PM is better for representing the evolution of WMH than IM when DEP-GAN is used, especially if ones would like to have good volumetric agreement and correlation, spatial agreement, and clinical plausibility of the WMH evolution. This is mostly due to false positives represented as changes observed in some cortical regions of the DEP model using IM due to brain atrophy and imaging artefacts.

### 4.3. Ablation study of the DEP-GAN’s regularisation terms

In this study, we proposed three regularisation terms for DEP-GAN (i.e., intensity, DSC, and volume) instead of one term (i.e., only intensity) like in the VA-GAN. Table 7 shows prediction results where the weights of each term are set to 0 to investigate how each of these three terms affect the prediction results. Note that λ_1_ is the weight for intensity loss, λ_2_ is the weight for DSC loss, and λ_3_ is the weight for volumetric loss (see Equation 4). We performed this ablation study using DEP-GAN-2C using PM. From this ablation study, the use of more terms in the regularisation had a positive impact in the prediction results. It is expected because multiple terms forced the DEP-GAN’s generator to generalise and perform well on all important measurements used in the evaluation of the prediction of WMH evolution, i.e., intensities in the regression of PM’s values, WMH segmentation correctness in DSC, and volumetric prediction of WMH. However, it is worth mentioning that the improvements were limited and still could be improved in the future.

**Table 7:** Results from ablation study of the DEP-GAN-2C’s regularisation terms tested using PM (see Equation 4). We calculated the prediction error of WMH change, volumetric agreement of WMH volume, and spatial agreement of WMH evolution, compared to the gold standard expert-delineated WMH masks (i.e., three-class DEM labels). “DSC” stands for similarity coefficient, “Int.” stands for intensity,“Vol.” stands for volumetric, “LoA” stands for limit of agreement, and “G” and “S” stand for percentage of subjects correctly predicted as having growing and shrinking WMH by DEP models. The best value for each learning approaches and evaluation measurements is written in bold.

| DEP-GAN-2C (PM) |  |  | Grow | Shrink | Avg. [%] | Vol. Bias [ml] | Lower | Upper | Entire | Change | Stable | Shrink | Grow | Avg. ((St+ |
| --- | --- | --- | --- | --- | --- | --- | --- | --- | --- | --- | --- | --- | --- | --- |
| $\lambda_1$ (Int.) | $\lambda_2$ (DSC) | $\lambda_3$ (Vol.) | (G) [%] | (S) [%] | ((G+S)/2) | mean(std) | LoA [ml] | LoA [ml] | WMH | (C) | (St) | (Sr) | (Gr) | Sr+Gr)/3) |
| 0 | 0 | 0 | 64.29 | 85.19 | **74.74** | 3.03(7.65) | -11.9684 | **18.0372** | 0.6131 | 0.1667 | 0.6178 | 0.1045 | 0.0813 | 0.2679 |
| 0 | 0 | 100 | 65.31 | 79.63 | 72.47 | 2.28(8.16) | -13.7197 | 18.2747 | **0.6132** | 0.1749 | 0.6166 | 0.1009 | 0.0909 | 0.2695 |
| 0 | 1 | 0 | 50.00 | 83.33 | 66.67 | 4.32(8.18) | -11.7181 | 20.3473 | 0.6093 | 0.1919 | 0.6063 | 0.1366 | 0.0706 | 0.2712 |
| 100 | 0 | 0 | 57.14 | 83.33 | 70.24 | 3.79(7.83) | -11.5525 | 19.1234 | 0.6075 | 0.1827 | 0.6143 | 0.1312 | 0.0741 | 0.2732 |
| 0 | 1 | 100 | **67.35** | 75.93 | 71.64 | 2.37(8.50) | -14.2904 | 19.0237 | 0.6101 | 0.1889 | 0.6177 | 0.1203 | 0.0922 | 0.2767 |
| 100 | 1 | 0 | 58.16 | 77.78 | 67.97 | **2.23(8.85)** | -15.1197 | 19.5748 | 0.6096 | 0.1912 | 0.6079 | 0.1209 | **0.0925** | 0.2738 |
| 100 | 0 | 100 | 57.14 | **88.89** | 73.02 | 4.51(8.15) | **-11.4546** | 20.4778 | 0.6078 | **0.1993** | 0.5996 | **0.1446** | 0.0760 | 0.2734 |
| 100 | 1 | 100 | 56.12 | 81.48 | 68.80 | 3.46(8.26) | -12.7218 | 19.6500 | 0.6107 | 0.1801 | **0.6245** | 0.1216 | 0.0868 | **0.2776** |

## 5. Conclusion and Future Work

In this study, we proposed a training scheme to predict the evolution of WMH using deep learning algorithms, namely Disease Evolution Predictor (DEP) model. We also proposed, evaluated, and studied different configurations of DEP models (i.e., with no human supervision or unsupervised (DEP-GAN using irregularity map), partial human supervision or semi-supervised (DEP-GAN using probability map), and full human supervision or fully supervised (DEP-UResNet) and different types of auxiliary input (i.e., Gaussian noise, WMH load, and WMH and stroke lesion loads) for prediction of WMH evolution. DEP models are more suitable for the problem of predicting WMH evolution than other models developed for predicting disease or stroke lesion evolution (Rekik et al., 2012, 2014) mainly because of two reasons: 1) DEP models use auxiliary input for modulating both image data (i.e., MRI) and non-image data (i.e., other risk factors) in different levels of convolutional layers and 2) DEP models can be configured to become probabilistic models and follow the nature of the prediction problem. To the best of our knowledge, this is the first extensive study on modelling WMH evolution using deep learning algorithms.

Based on the two ablation analyses done as part of the present study, DEP-GAN with 2 critics (DEP-GAN-2C) performed better than WGAN-GP, VA-GAN, and DEP-GAN using 1 critic (DEP-GAN-1C). We would like to emphasise the importance of the two critics in our proposed DEP-GAN model. While it is possible to perform direct regression of DEM values and not using GAN (as we have “paired” baseline and follow-up images), our previous experiments indicate that the direct regression using deep neural networks without critics did not work properly for our task and its performance was worse than our proposed DEP-GAN model. We observed that the changes of IM/PM values from baseline to follow-up (i.e., the DEM values), in addition to their subtlety, are small or very small, resulting in very sparse data. This causes the deep neural network (i.e., the generator) to struggle, failing to learn anything useful from the training data when regressing IM/PM values. Thus, it produced only zeros to all voxels in the testing stage despite the mean squared error (MSE) value being very low and close to zero in the training. This is a strong indication of strong overfitting when no discriminators/critics are used, i.e. with coefficients equal zero or close to zero, the terms in the regression model also become zero, and the fitness expressed in the MSE rather than reflecting a true model fit expresses the fitness of a model that barely has any meaningful terms. By using two critics, the generator has to learn not only the regression of DEM values but also the “context” of the DEM while performing realistic modifications to the follow-up images.

Furthermore, Gaussian noise successfully improved all DEP models in almost all evaluation measurements when it was used as auxiliary input, probably because it compensates for the sparsity of the input data. At the same time, it shows that there are indeed some unknown factors that influence the evolution of WMH. These unknown factors make the problem of predicting/delineating WMH evolution non-deterministic, and Gaussian noise were proposed to simulate this scenario. The intuition behind this approach is that Gaussian noise fills in the missing (unavailable) risks factors or their combination, which could influence the evolution of WMH. Note that it is very challenging to collect and compile all risk factors of WMH evolution in a longitudinal study.

From our experiments, on average, DEP-UResNet (i.e., a fully supervised scheme) yielded the best results in almost every evaluation measurement. However, it is worth to mention that it did not perform well in the clinical plausibility test. DEP-GAN-2C using PM yielded similar average performance to the DEP-UResNet’s performance and yielded the best results out of all schemes in the clinical plausibility test. Moreover, results from DEP-UResNet and DEP-GAN-2C using PM were not statistically different to each other on delineating the WMH clusters.

If we consider the results, time, and resources spent in this study, then DEP-GAN-2C using PM showed the biggest and strongest potential of all DEP models. Not only did it perform similarly to the DEP-UResNet but it did not need manual WMH labels in baseline and follow-up scans for training. The PM needed as input for this model can be efficiently produced by any supervised deep/machine learning model. Moreover, the development of automatic WMH segmentation for producing better PM could be done separately and independent from the development of the DEP model. If a better PM model is available in the future, then the model can be retrained using the newly produced PM for better performance. Also, DEP-GAN-2C using PM could be used for other (neurodegenerative) pathologies, as long as a set of PM from these other pathologies is produced and used to (re-)train the model.

There are several shortcomings anticipated from the results of this study. Firstly, manual WMH labels of two MRI scans (i.e., baseline and follow-up scans) are necessary for training the DEP-UResNet. In many scenarios, this is not practical and efficient in terms of time and resources. Secondly, the DEP-GAN-2C using IM is computationally very demanding as it involves regressing IM values across the whole brain tissue. This resulted in low performances in almost all evaluation measurements. Thirdly, the schemes’ performances depend on the accuracy of the quality of input. For example, the PM generated in this study are slightly biased towards overestimating the WMH in the optical radiation and underestimating WMH in the frontal lobe. This could be caused by the absence of correcting the FLAIR images for b1 magnetic field inhomogeneities. However, a previous study on small vessel disease images demonstrated this procedure might affect the results underestimating the subtle white matter abnormalities characteristics of this disease, and recommends this procedure to be used in T1- and T2-weighted structural images but not in FLAIR images for WMH segmentation tasks (Hernández et al., 2016). Hence, the biggest challenge of using DEP-GAN-2C using PM is its highly dependency on the quality of initial PM. Fourthly, volumetric agreement analyses suggest that there are still large differences in absolute volume and in change estimates produced by the proposed DEP models. While this study is intended as a “proof-of-principle” study to advance the field of white matter -and ultimately brain health-prediction, it is worth to mention that better reliability in the WMH assessment is necessary so as DEP models can be used in clinical practice. Furthermore, better understanding of what DEP models extract to estimate WMH evolution would be very useful in clinical practice. Lastly, the limitation of using (Gaussian) random noise in DEP models is the fact that we do not really know which set of Gaussian random noise should be used to generate the best result for each subject. Note that, in this study, all DEP models that used Gaussian noise as auxiliary input were tested 10 times to calculate the mean and the “best” set of Gaussian noise which produced the best automatic delineation of WMH evolution overall. In conclusion, DEP models suffer similar problems and limitations to any machine learning based medical image analysis methods.

The DEP models proposed in this study open up several possible future avenues to further improve their performances. Firstly, multi-channel (e.g., PM and T2-FLAIR) input could be used instead of single channel input. In this study, we only used single channel to draw a fair comparison between DEP-UResNet which uses T2-FLAIR and DEP-GAN which uses either IM or PM. Secondly, 3D architecture of DEP-GAN could be employed when more subjects are accessible in the future. 3D deep neural networks have been reported to have better performances than the 2D ones, but they are more difficult to train (Çiçek et al., 2016; Baumgartner et al., 2017). Thirdly, enhanced learning techniques such as transfer learning and advance data augmentation can be applied in future studies to improve the performance of DEP models. Fourthly, Gaussian noise and known risk factors (e.g., WMH and SL loads) could be modulated together instead of modulating them separately in different models. By modulating them together, the DEP model would be influenced by both known (available) risk factors and unknown (missing) factors represented by Gaussian noise. Lastly, different random noise distribution could be used for auxiliary input. Note that each risk factor of WMH evolution (e.g., WMH load, age, and blood pressure) could have different data distribution, not only Gaussian distribution with zero mean and unit variance. If a specific data distribution (i.e., the same or similar to the real risk factor’s data distribution) could be used for a specific risk factor, then the real data could replace the random noise if available in the testing.

## Acknowledgements

Funds from the Indonesia Endowment Fund for Education (LPDP), Ministry of Finance, Republic of Indonesia (MFR); Row Fogo Charitable Trust (Grant No. BRO-D.FID3668413)(MCVH); Wellcome Trust (patient recruitment, scanning, primary study Ref No. WT088134/Z/09/A); Fondation Leducq (Perivascular Spaces Transatlantic Network of Excellence); EU Horizon 2020 (SVDs@Target); and the MRC UK Dementia Research Institute at the University of Edinburgh (Wardlaw programme) are gratefully acknowledged. The Titan Xp used for this research was donated by the NVIDIA Corporation.

## Appendix A. Volumetric agreement and correlation graphs

**Figure A.10:**
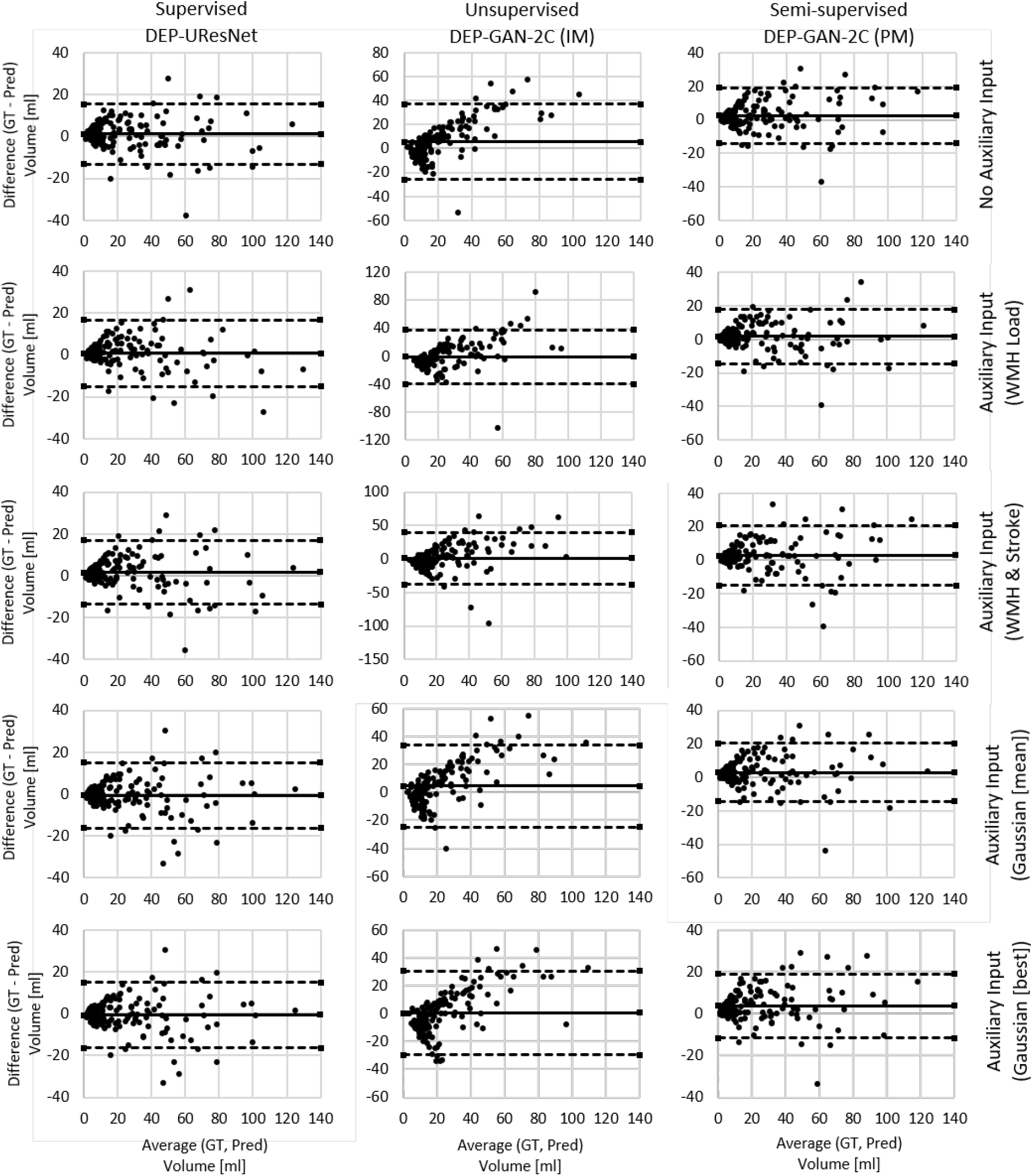
Volumetric agreement analysis (in *ml*) between ground truth (GT) and predicted volume of WMH with different types/modalities of auxiliary input (Pred) using Bland-Altman plot which correspond to data presented in Table 4. Solid lines correspond to “Vol. Bias” while dashed lines correspond to either “Lower LoA” or “Upper LoA” of the same table. “LoA” stands for limit of agreement.

**Figure A.11:**
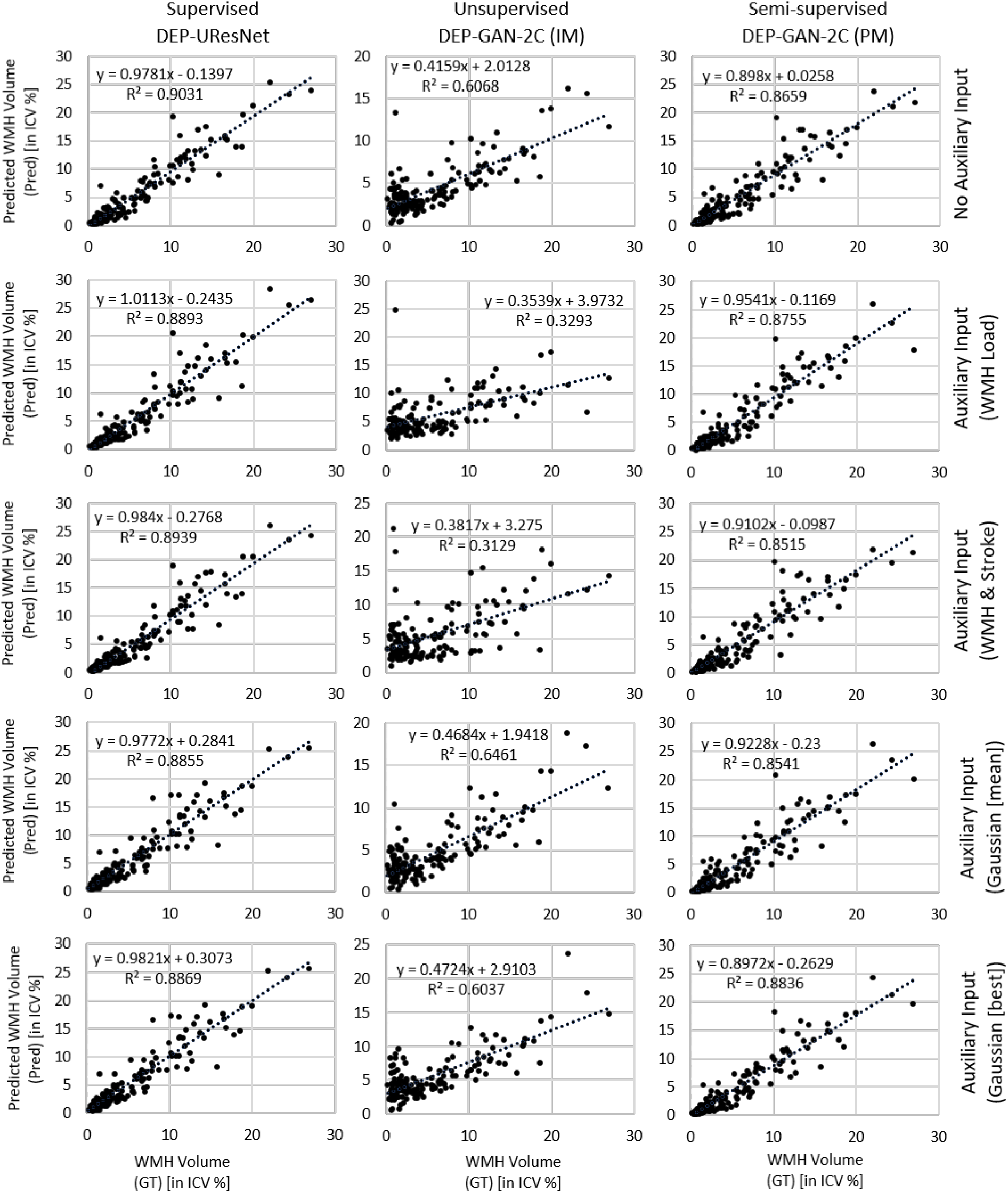
Correlation plots between manual WMH volume produced by the expert (GT) and predicted WMH volume by various DEP models with different types/modalities of auxiliary input (Pred). WMH volume is in the percentage of intracranial volume (ICV) to remove any potential bias associated with head size.

